# Hyperosmotic Stress of HL-60/S4 Cells: Gene Set Enrichment Analysis and Microscopic Changes

**DOI:** 10.1101/2025.04.22.649232

**Authors:** Ada L. Olins, David Mark Welch, Igor Prudovsky, Donald E. Olins

## Abstract

Cellular homeostasis requires that internal cell conditions (e.g., pH, ionic and non-ionic solute concentrations and hydration levels) maintain a dynamic stability. To better understand the mechanisms enforcing this homeostasis, we examined the transcript abundance and cellular morphology of undifferentiated myeloid HL-60/S4 cells exposed to 30 or 60 minutes of hyperosmotic stress, resulting from 300 mM sucrose added to the tissue culture medium. We used Gene Set Expression Analyses (GSEA) to examine various cell functions (Phenotypes) of interest. Microscopy was also employed, where possible, to visualize the nuclear and chromatin physical states. We observed that hyperosmotic stress resulted in congelation of interphase and mitotic chromatin. In parallel, GSEA indicated a reduction of heterochromatin, histone lysine methyltransferase and mRNA transcription. Mitosis ceases during hyperosmotic stress, with the intact tubulin spindle remaining attached to clustered centromeres on the congealed chromosomes. GSEA indicated that ribosome biosynthesis in nucleoli and oxidative activity in mitochondria are significantly increased. In addition, the proteasome phenotype is increased, suggesting that protein synthesis and destruction are both occurring at an increased pace during hyperosmotic stress. Nuclear envelope-associated chromatin also appears to be affected by hyperosmotic stress: LBR and Heterochromatin Protein 1 alpha (CBX5) disperse into the cytoplasm. Evidence is also presented that 300 mM sucrose leads to a reduction of DNA methylation and aberrant cellular localization of MeCP2. The effects of acute hyperosmotic stress on HL-60/S4 cells are very diverse and very profound. In many respects, the stress response resembles a frantic attempt for cell survival in the face of inevitable cell death.

## Introduction

Cellular hyperosmotic stress, by external cell membrane-impermeable non-ionic or ionic dissolved solutes, leads to cell shrinkage and molecular crowding. This results in nuclear chromatin congelation (condensation) and phase separation with adverse effects upon numerous cellular functions (Finan and Guilak, 2010; Finan et al., 2011; Irianto et al., 2013; Mark Welch et al., 2022; Olins et al., 2020; Richter et al., 2007). In our most recent article on HL-60/S4 cell functional changes resulting from sucrose induced hyperosmotic stress (Mark Welch et al., 2022), we employed Gene Ontology (GO) analyses of the transcription data to identify some of the perturbed functions. The present study is an extension of these GO analyses, by employing Gene Set Enrichment Analysis (Mootha et al., 2003; Subramanian et al., 2005), which identifies functional cellular states (Phenotypes) that exhibit “enrichment” of diagnostic gene sets for any particular Phenotype. This present analysis is based upon differential gene expression (DGE), comparing undifferentiated HL-60/S4 cells exposed to 300 mM sucrose (added to tissue culture medium) for 30 or 60 minutes, with cells in tissue culture medium without sucrose. In addition, we report results of immunostaining and FISH (Fluorescent In Situ Hybridization) microscopy, employed to visualize chromatin and nuclear structural changes that parallel the GSEA functional interpretations.

## Materials and Methods

### Cell Cultivation

HL-60/S4 cells were cultivated in the following medium: RPMI 1640 + 10% (unheated) Fetal calf serum + 1% Pen/Strep/Glut. Generally, 5 ml of growing HL-60/S4 cells were added to a T-25 flask containing dry sucrose to yield ∼10^6^ cells/ml in a medium of ∼600 mOsM. Further treatment of the cells has been described earlier (Mark Welch et al., 2022; Olins et al., 2020). Cell death became apparent by 12-24 hours; experiments only continued for 30 and 60 minutes.

### Gene Set Enrichment Analysis (GSEA)

The transcriptome data of HL-60/S4 (+/- sucrose) can be found in “Table S1 Transcriptome Data.xlsx”. We use GSEA (https://www.gsea-msigdb.org/gsea/login.jsp) to match an experimental differential gene expression list “Table S2 Formatted for Expression dataset.txt” against gene sets from the Molecular Signature Database (MSigDB), which contains functionally related genes (Mootha et al., 2003; Subramanian et al., 2005). GSEA software ranks the genes in the gene set according to the experimental level of expression. Analysis of this ranked gene list documents the differential gene expression between two cell phenotypes. The Enrichment Score (ES) can be ES > 0.00 or < 0.00, reflecting the extent to which the gene set is overrepresented (+, upregulated) or underrepresented (-, downregulated). The nominal p-value is an estimate of the statistical significance of an ES, where p = 0.00 is the maximal significance; p > 0.05 is regarded as less significant. The GSEA User Guide section on “Nominal P Value” describes that p-value accuracy can be increased by increasing permutations during the calculations (e.g., 100 to 1000 or more permutations). In most cases, we have employed 1000 permutations. Normalized Enrichment Score (NES) permits comparisons between different GSEA analyses, by accounting for variations in different gene set sizes. An important additional feature of GSEA is identification of the “Leading Edge”, a subset (core) of the ranked gene set that contributes most to the ES. Analysis of the Leading Edge is a useful method for identification of the most influential genes in establishing a phenotype, which we have utilized quite frequently.

The GSEA website (https://www.gseamsigdb.org/gsea/index.jsp) explains how to conduct analyses and lists many functionally defined gene sets. Gene sets used were chosen from the Molecular Signature Database (MSigDB). We compared all GO gene sets from MSigDB, selecting the “Top 10” and “Top 20” gene sets with nominal p values < 0.05 and beginning at highest NES values. The gene sets from “Top 10” were selected to compare Ribosome Biogenesis (Table 4); from “Top 20”, to compare Mitochondrial Function (Table 5).

It is important to point out that GSEA is inherently “predictive” of the cellular phenotypes, derived by analyses of the experimental relative gene expression data. The gene sets assigned to particular phenotypes assume that the mRNAs are translated into proteins reflecting their relative gene expression. Verification of these predictions would require extensive quantitative protein analysis.

### Microscopy and Immunostaining

Leica SP8 confocal imaging of HL-60/S4 cells (+/- 300 mM sucrose) employed the following antibodies (including their source and dilution):

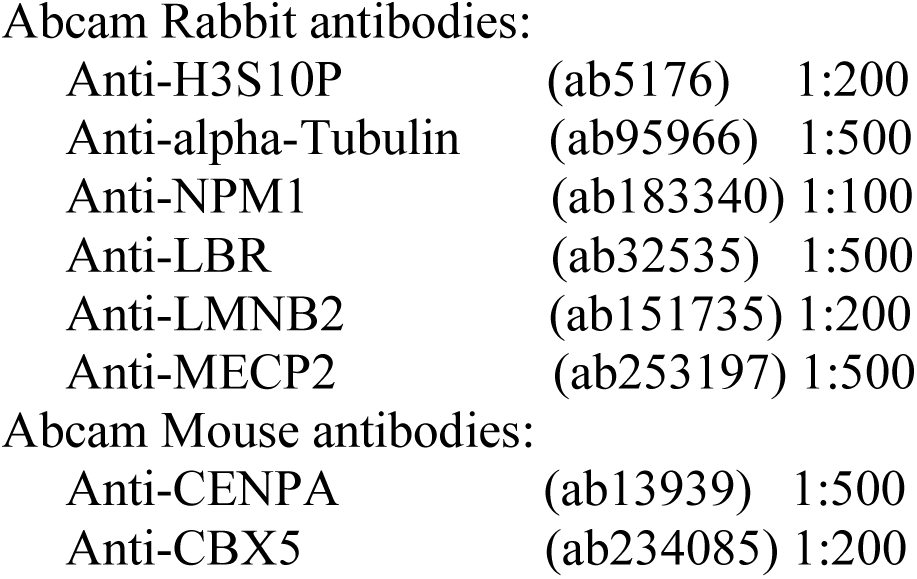

|  |  |  |
| --- | --- | --- |
| Anti-H3S10P | (ab5176) | 1:200 |
| Anti-alpha-Tubulin | (ab95966) | 1:500 |
| Anti-NPM1 | (ab183340) | 1:100 |
| Anti-LBR | (ab32535) | 1:500 |
| Anti-LMNB2 | (ab151735) | 1:200 |
| Anti-MECP2 | (ab253197) | 1:500 |
| Abcam Mouse antibodies: |  |  |
| Anti-CENPA | (ab13939) | 1:500 |
| Anti-CBX5 | (ab234085) | 1:200 |

*In Vivo* staining of mitochondria activity: MitoTracker Red CMXRos was purchased from Cell Signaling. The protocol employed is as follows: Prepare a 1 mM stock solution of MitoTracker in DMSO. Add 3μl of stock solution to 10 ml of cells in medium. Split 5 ml of cells into a T-25 flask with dry sucrose to make cells in 300 mM sucrose; keep 5 ml in flask for control (0).

Incubate both 0- and 300-mM sucrose flasks for 30 minutes at 37° C. Centrifuge cells. Wash the separate cell pellets in the appropriate PBS (+/-) sucrose. Load cell suspensions onto fresh polylysine-coated slides. Let cells settle for 30 minutes. Drain slides and fix with 3.7% formaldehyde. Quench the formaldehyde with 50 mM NH4Cl for 1 minute. Wash slides in PBS. Permeabilize the attached fixed cells with Triton X-100/PBS for 20 minutes. Wash slides in PBS. Mount coverslips in Vectashield+DAPI.

RNA stain: Syto RNASelect was purchased from ThermoFisher Scientific and tested following their protocol. All micrographs presented are single slices from deconvolved confocal stacks.

## Results

### Hyperosmotic Stress with Sucrose produces Congealed Mitotic Chromosomes and Congealed Interphase Chromatin

In our first article on the structural effects of acute hyperosmotic stress upon *in vivo* chromatin (Olins et al., 2020), we employed microscopic imaging of DAPI (DNA) stained undifferentiated HL-60/S4 cells. The cells were incubated in tissue culture medium with-or-without 300 mM sucrose for 30 or 60 minutes. Mitotic chromosomes collapsed into an amorphous chromatin aggregate with the epigenetic mitotic “marker” (i.e., phosphorylated H3S10, “H3S10P”) segregated at the surface of the congealed chromosomes (Figure 1). The “fine” interphase chromatin appears to form much thicker fibers in the hyperosmotic conditions (Figure 2). In addition, we previously published numerous examples of “phase separation” of chromatin-binding proteins, demonstrated by immunostaining (e.g., Ki67, CTCF, RAD21, HMGN2, HMGB2, H1.2 and H1.5), following exposure of the live HL-60/S4 cells to acute hyperosmotic stress, see (Olins et al., 2020).

**Figure 1.**
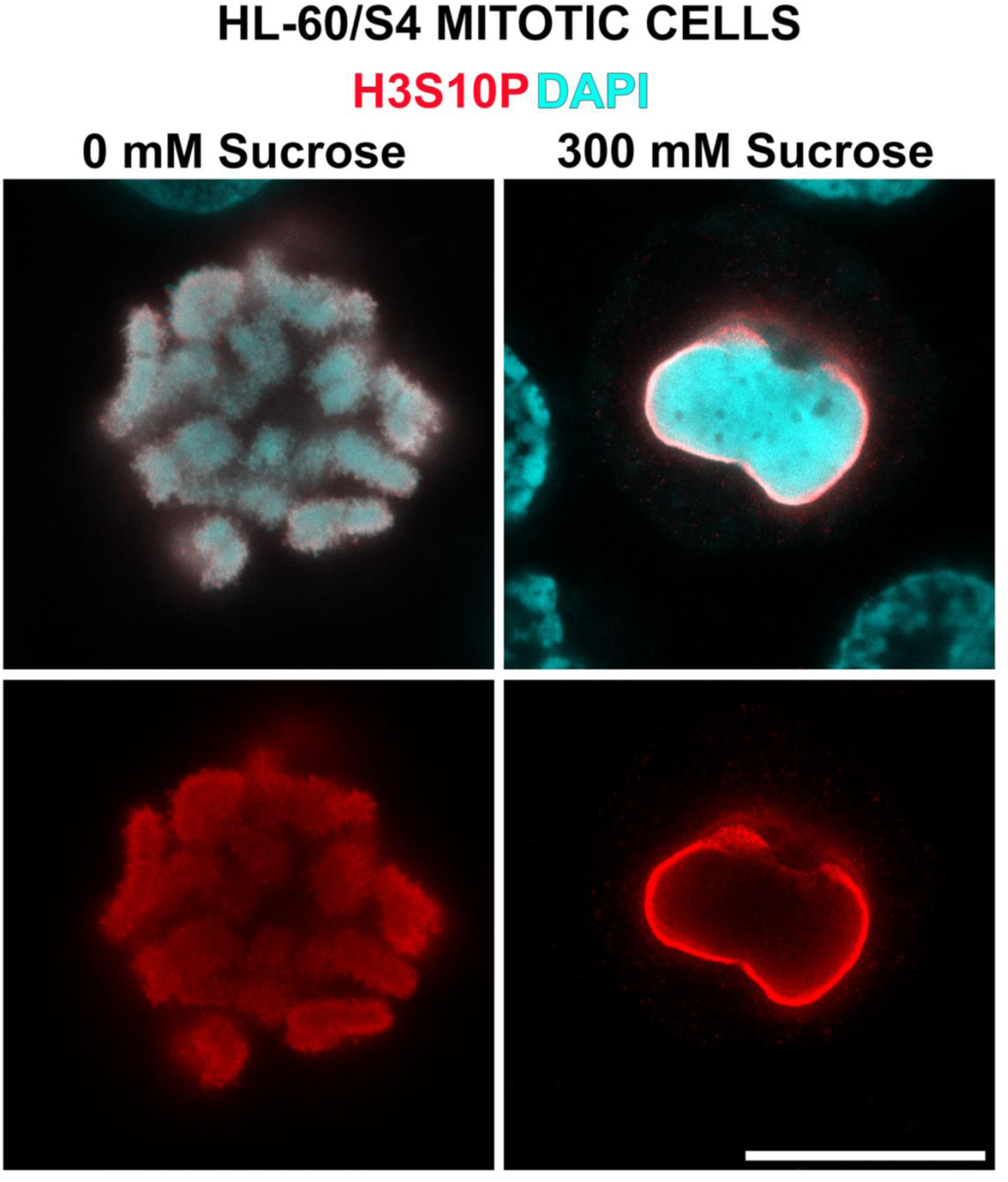
Effect of acute hyperosmotic stress upon mitotic chromosome DNA distribution and localization of the mitotic chromosome “marker” (H3S10P). Presented are confocal slices of undifferentiated HL-60/S4 cells after fixation, permeabilization and immunostaining. Upper left panel: Control cell mitotic chromosomes <u>untreated</u> in tissue culture medium. Upper right panel: Congealed mitotic chromosomes in tissue culture medium plus 300 mM sucrose for 30 minutes (∼600 mOsM). The Bottom panels display only H3S10P localization in the corresponding Upper panels. Colors: H3S10P (red); DAPI (cyan). Magnification bar: 10 μm. Similar images can be found in (Olins et al., 2020).

**Figure 2.**
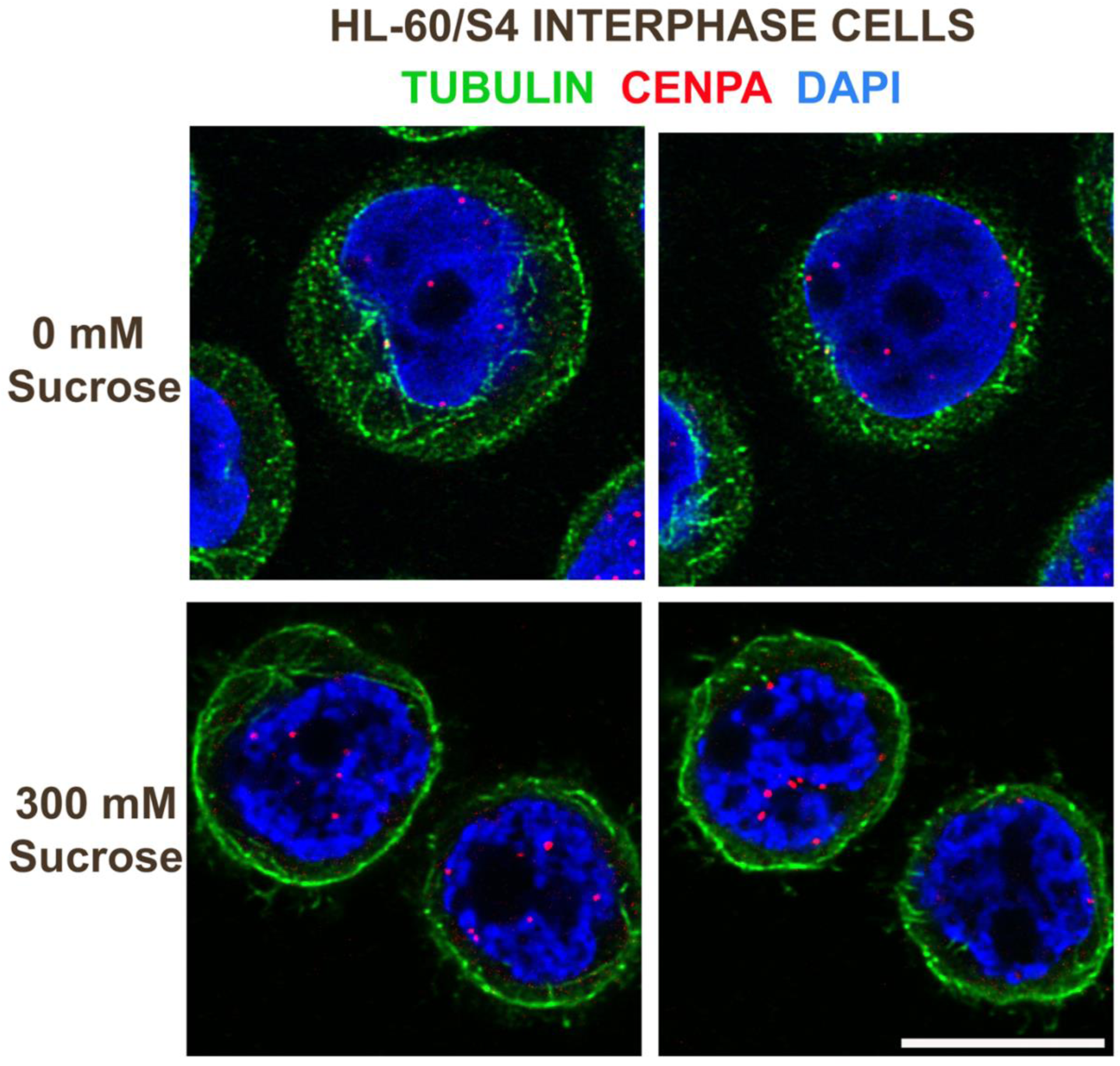
Effect of acute hyperosmotic stress (∼600 mOsM) upon interphase chromatin condensation and distribution (DAPI). Two slices from the deconvolved image stacks are shown for each osmolarity level. Besides the chromatin congelation in 300 mM sucrose, cytoplasmic alpha-tubulin (green filaments) concentrates at the plasma membrane. The centromere marker CENPA (red dots) appears to localize and cluster at the surface of the congealed chromatin fibers. Magnification bar: 10 μm.

### Despite the Congelation of Interphase Chromatin by Hyperosmotic Stress, Gene Set Enrichment Analysis (GSEA) indicates reduced transcript levels for genes associated with Heterochromatin Formation

Figure 3 presents three panels of Heterochromatin related GSEA enrichment plots for 30 minutes stress *versus* 0 minutes control. Table 1 presents GSEA parameters for heterochromatin enrichment/depletion, comparing stressed undifferentiated HL-60/S4 cells to unstressed undifferentiated control cells. It is clear that, for these three heterochromatin panels, there are significant decreases in NES (Normalized Enrichment Score). This result implies an actual “reduction” in heterochromatin-related transcripts. The anti-correlation between “increased” chromatin congelation and “decreased” heterochromatin suggests that congealed chromatin and heterochromatin may have very different structures. The fourth panel in Figure 3 (GOBP_MRNA_TRANSCRIPTION) illustrates that mRNA levels for genes involved in mRNA transcription are reduced following 30 minutes of sucrose stress. This observation should be compared to a previous analysis: Table 1 of (Mark Welch et al., 2022), which demonstrated that ∼21% of the total mapped genes (∼16,000 genes) are downregulated, ∼18% are upregulated and ∼80% unchanged after 30 minutes of sucrose stress. Table 1 of (Mark Welch et al., 2022) is not restricted to genes involved in mRNA synthesis, as is the fourth panel in Figure 3.

**Figure 3.**
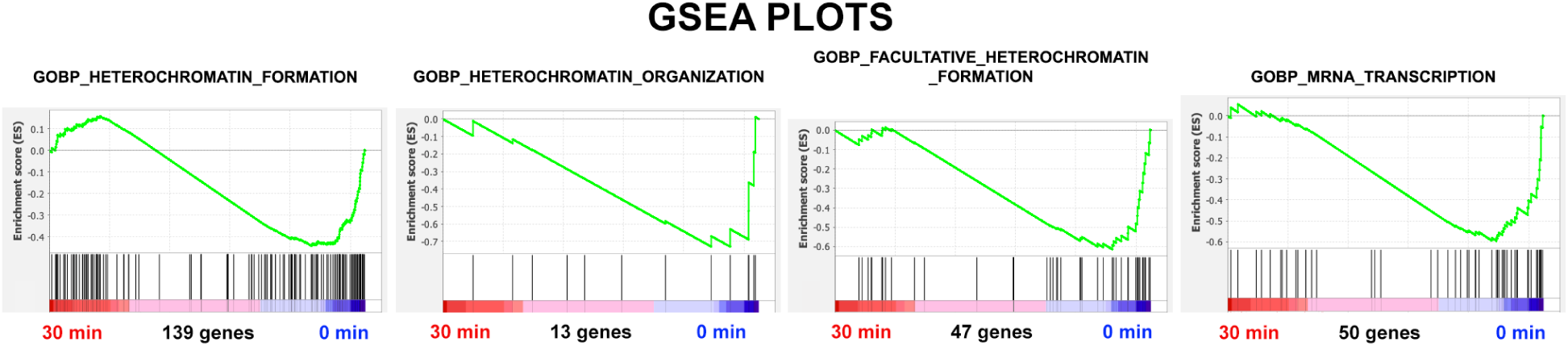
Three GSEA enrichment plots of Heterochromatin related transcripts, demonstrating reduction of NES after 30 minutes of exposure to Hyperosmotic stress conditions, compared to unstressed undifferentiated HL-60/S4 cells control (0) cells. The fourth GSEA enrichment plot is concerned with the transcription of a specific gene set involved with mRNA transcription (e.g., TATA-box binding proteins), displaying a reduction of NES, comparing 30 minutes of hyperosmotic sucrose to unstressed undifferentiated HL-60/S4 cells control cells.

**Table 1.**
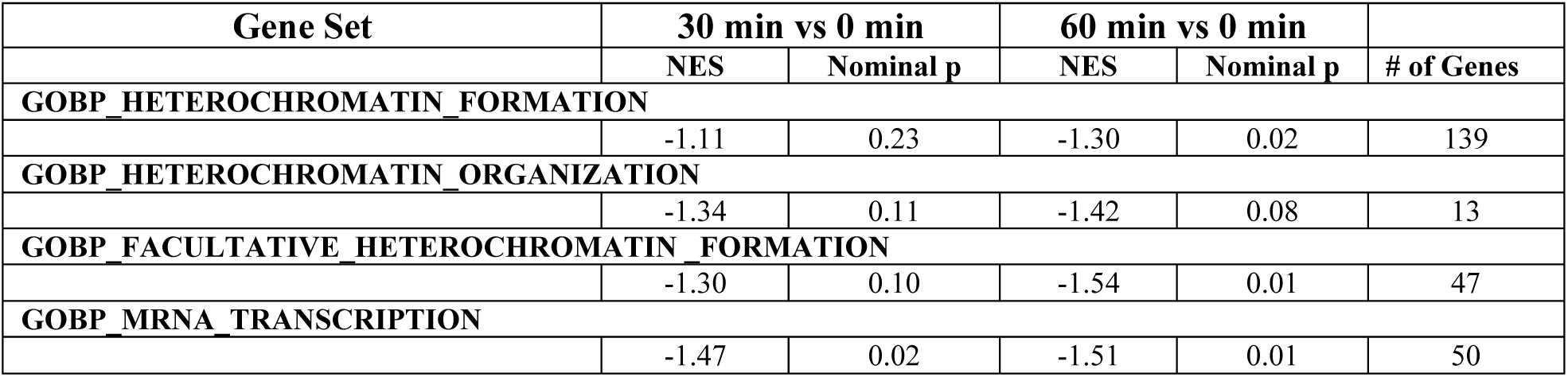
GSEA parameters for Heterochromatin Formation and for mRNA transcription, comparing 30 and 60 minutes stressed undifferentiated HL-60/S4 cells to unstressed control cells: NES, Normalized Enrichment Score; Nominal p, statistical significance, where p = 0.00 is the maximal significance and p > 0.05 is regarded as less significant.

| Gene Set | 30 min vs 0 min |  | 60 min vs 0 min |  | # of Genes |
| --- | --- | --- | --- | --- | --- |
|  | NES | Nominal p | NES | Nominal p |  |
| GOBP_HETEROCHROMATIN_FORMATION |  |  |  |  |  |
|  | -1.11 | 0.23 | -1.30 | 0.02 | 139 |
| GOBP_HETEROCHROMATIN_ORGANIZATION |  |  |  |  |  |
|  | -1.34 | 0.11 | -1.42 | 0.08 | 13 |
| GOBP_FACULTATIVE_HETEROCHROMATIN_FORMATION |  |  |  |  |  |
|  | -1.30 | 0.10 | -1.54 | 0.01 | 47 |
| GOBP_MRNA_TRANSCRIPTION |  |  |  |  |  |
|  | -1.47 | 0.02 | -1.51 | 0.01 | 50 |

Leading Edge analysis of the three Heterochromatin GSEA enrichment plots shown above (Figure 3 and Table 1) revealed a large number of common downregulated genes. For example, the entire “Facultative” Leading Edge is included within the larger “Heterochromatin Formation” Leading Edge. Strikingly, this includes histone lysine methylation and demethylation transcripts (e.g., KMT2A and KDM5A). Their combined downregulation suggests that this form of epigenetic modification is “slowed-down” by the hyperosmotic stress.

### Hyperosmotic Stress results in a decreased mRNA levels for Histone Lysine Methyltransferase enzymes

Figure 4 and Table 2 present a GSEA comparison of mRNA levels for histone methyltransferases. Focusing on these enzymes is consistent with the prior conclusions that hyperosmotic stress is responsible for the diminishing heterochromatin.

**Figure 4.**
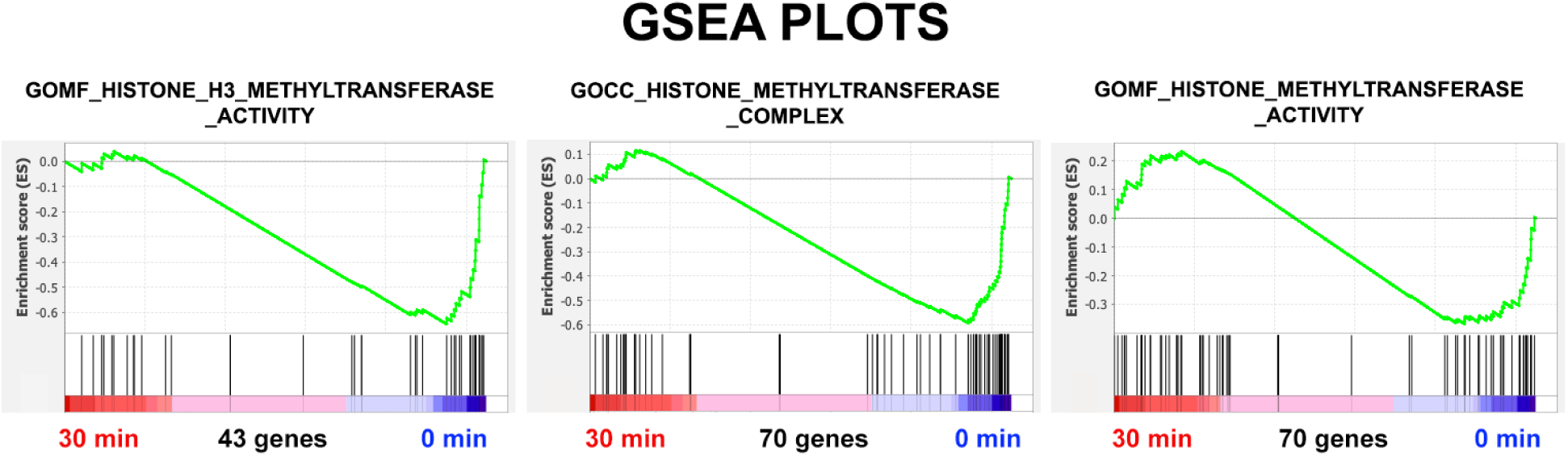
Three GSEA enrichment plots of Histone Methyltransferase transcripts, demonstrating reduction of NES after 30 minutes of exposure to Hyperosmotic stress conditions, compared to unstressed undifferentiated HL-60/S4 cells control (0) cells.

**Table 2.** GSEA parameters for Histone Methyltransferase transcripts, comparing 30 and 60 minutes stressed undifferentiated HL-60/S4 cells to unstressed control cells: NES, Normalized Enrichment Score; Nominal p, statistical significance, where p = 0.00 is the maximal significance and p > 0.05 is regarded as less significant.

| Gene Set | 30 min vs 0 min |  | 60 min vs 0 min |  | # of Genes |
| --- | --- | --- | --- | --- | --- |
|  | NES | Nominal p | NES | Nominal p |  |
| GOMF_HISTONE_H3_METHYLTRANSFERASE_ACTIVITY |  |  |  |  |  |
|  | -1.39 | 0.04 | -1.63 | 0.0 | 43 |
| GOCC_HISTONE_METHYLTRANSFERASE_COMPLEX |  |  |  |  |  |
|  | -1.32 | 0.05 | -1.63 | 0.0 | 70 |
| GOMF_HISTONE_METHYLTRANSFERASE_ACTIVITY |  |  |  |  |  |
|  | -0.99 | 0.47 | -1.16 | 0.18 | 70 |

In a similar fashion to the prior analysis of Heterochromatin transcripts, Leading Edge analysis of the three methyltransferase enrichment plots shown above (Figure 4 and Table 2) revealed a large number of common downregulated genes “Table S3 Methyltransferase Leading Edges.xlsx”. GOMF_HISTONE_H3_METHYLTRANSFERASE _ACTIVITY and GOMF_HISTONE_METHYLTRANSFERASE _ACTIVITY Leading Edges were essentially identical (except the former gene list, which was missing one out of 20 genes, KMT5A). GOCC_HISTONE_METHYLTRANSFERASE _COMPLEX has 32 genes, of which 24 are different than the other Leading Edges, including decreases in the TATA-Box binding proteins (TAF1 and TAF4).

### Hyperosmotic Stress with Sucrose appears to block Mitosis with the Microtubular Spindle Apparatus remaining intact and continuing to adhere to the congealed Mitotic Chromosome Centromeres

In addition to documenting the Hyperosmotic Stress-induced interphase chromatin congelation Figure 2), Figure 5 demonstrates the stability of the attachment of spindle microtubules to the congealed mitotic chromosome centromeric regions, highlighted by immunostaining with anti-CENPA. Whereas, the stained centromeres are scattered around the unstressed mitotic chromosomes (Figure 5, top row), they tend to cluster at the surface of the stressed congealed mitotic chromosomes (Figure 5, bottom two rows).

**Figure 5.**
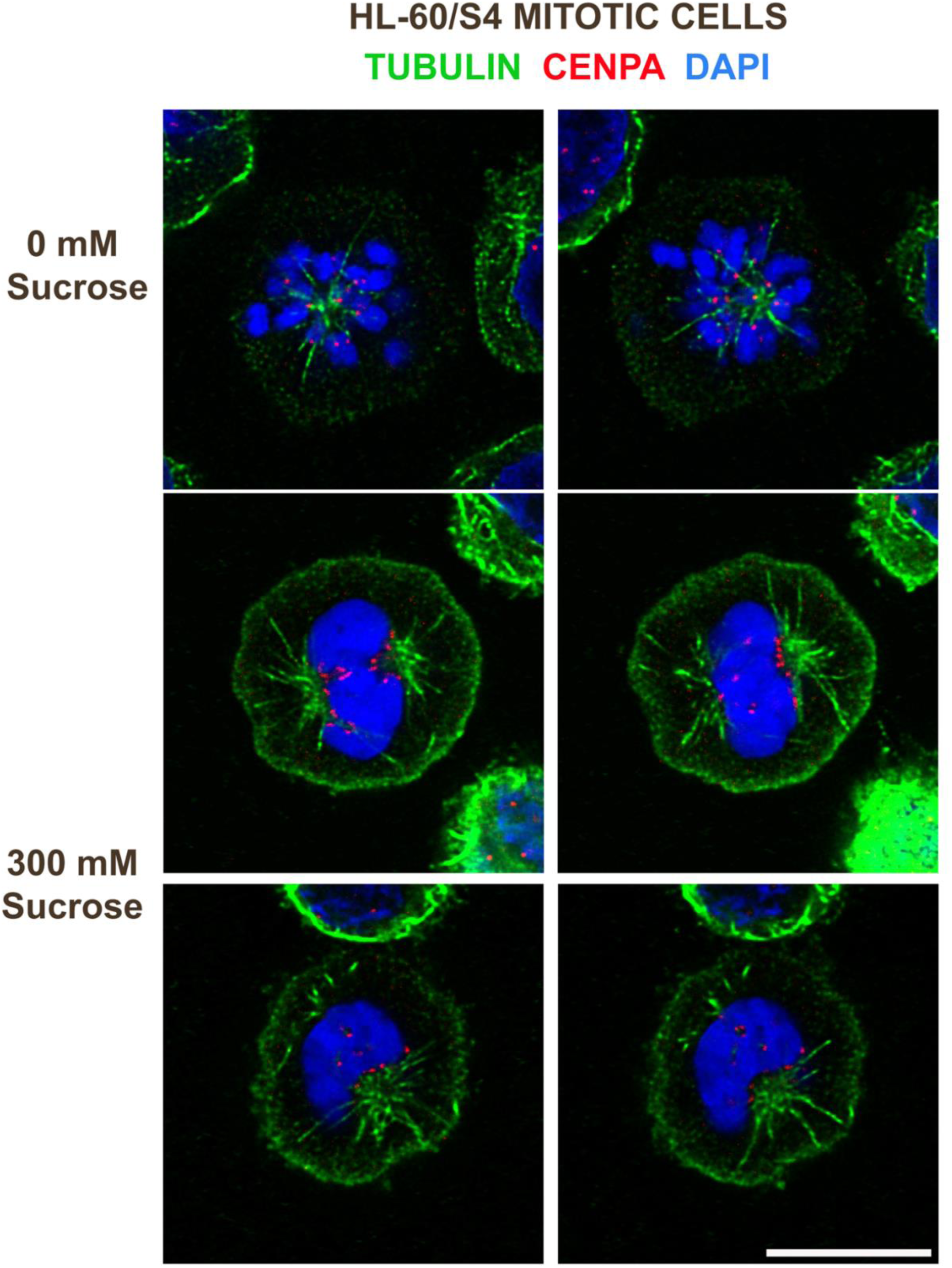
Two slices from the deconvolved image stacks of three mitotic cells without (top row) or with (bottom two rows) incubation for 30 minutes in 300 mM sucrose added to tissue culture medium. Note that the spindle microtubules (green) remain attached to the exposed centromeres (CENPA, red) even in the congealed mitotic chromosome clusters. In addition, many cytoplasmic microtubules exhibit proximity to the (presumptive) plasma membrane. Magnification bar: 10 μm

### GSEA plots demonstrate that Hyperosmotic Stress with Sucrose yields considerable enrichment of transcripts related to Centromere Structure and Function

In an effort to understand which gene transcripts might be most important in maintaining the microtubule-centromere complex during hyperosmotic conditions, we examined the Leading Edges of the four gene sets displayed in Figure 6 and Table 3. Employing a Venn Diagram, we identified the most prevalent common genes in the four Leading Edges. Not surprisingly, these genes included major structural components of the centromeres (e.g., CENP proteins) and numerous histone H4 variants. Earlier studies (Tanaka and Desai, 2008) have indicated that other proteins (i.e., NDC80, NUF2, BCAS2 and AURKB) are involved in the attachment of microtubules to the kinetochore. Analysis of Log2FC (Fold Changes) of the transcriptomes of undifferentiated HL-60/S4 cells exposed to 300 mM sucrose for 30 and 60 minutes indicates that these (and other) gene transcripts are elevated by the hyperosmotic stress conditions (Figure 7). Clearly, numerous factors cooperate to stabilize the mitotic complex during hyperosmotic stress.

**Figure 6.**
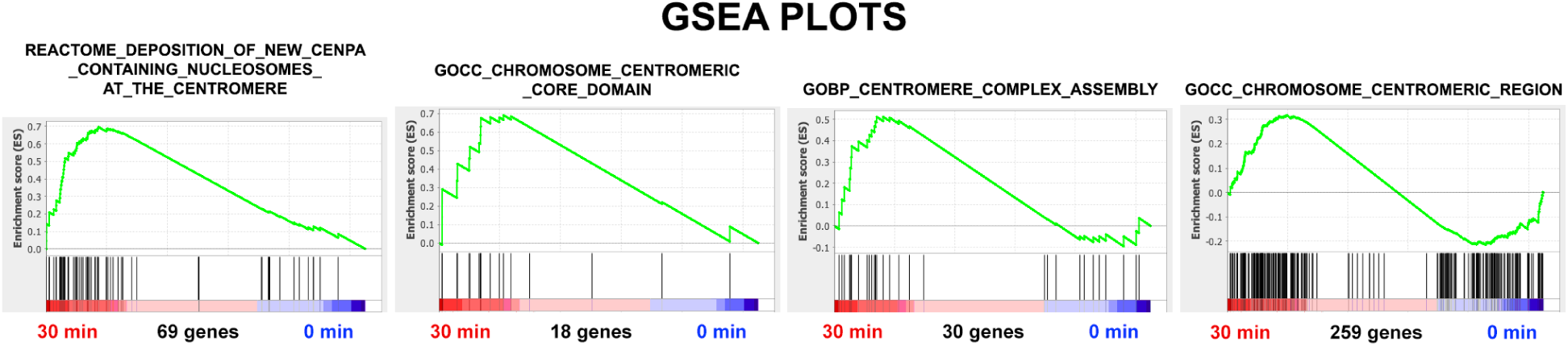
GSEA enrichment plots of Centromere-related transcripts demonstrating enrichment after 30 minutes of exposure to Hyperosmotic stress conditions, comparing stressed (30 min) undifferentiated HL-60/S4 cells to unstressed control (0 min) cells.

**Figure 7.**
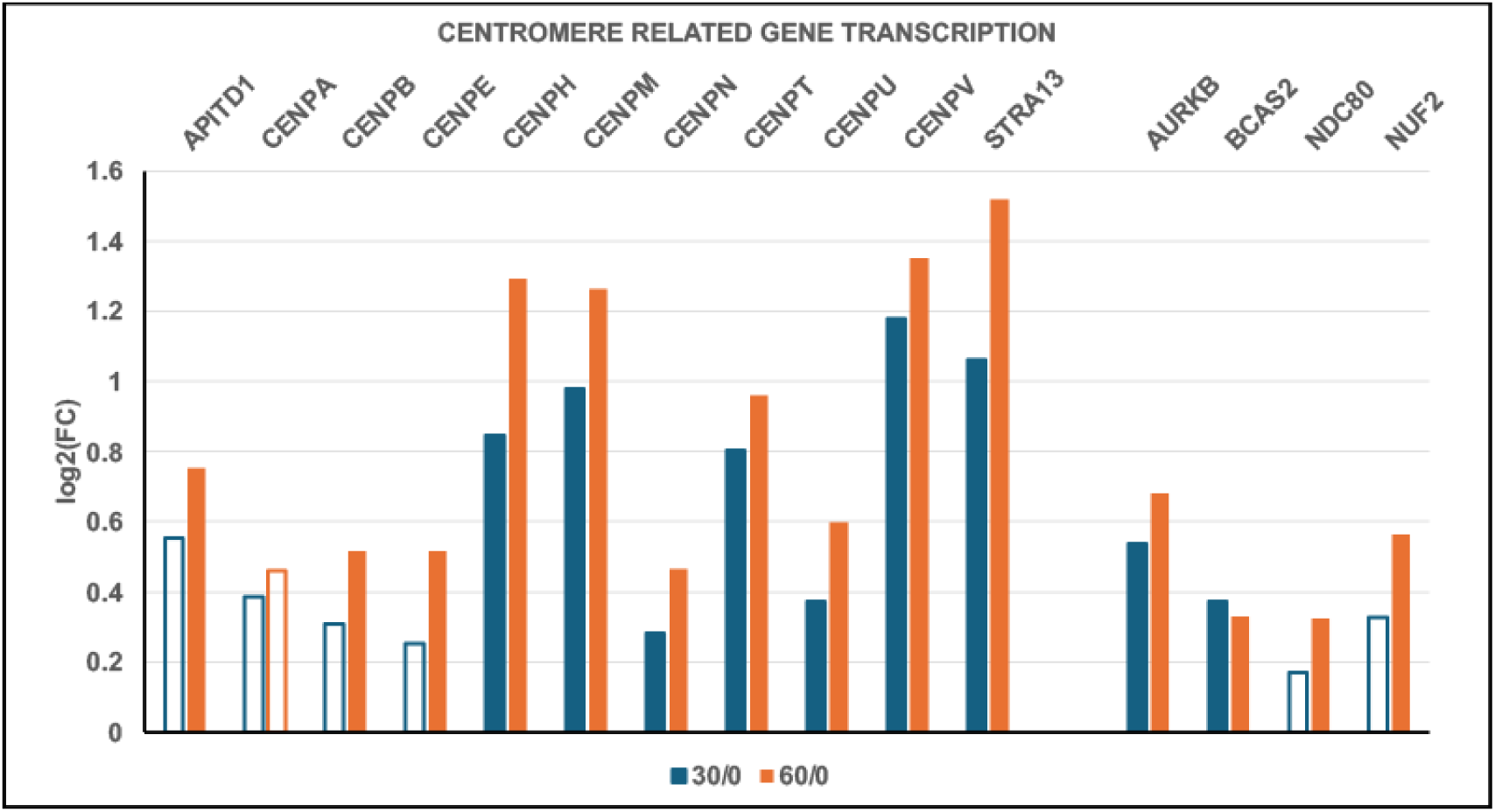
Log2FC Gene Transcript changes of Centromere Structural Proteins and other proteins involved in Attachment of Microtubules to the Kinetochore, comparing 30 and 60 minutes of Hyperosmotic Stress to Unstressed (0) undifferentiated HL-60/S4 Cells (i.e., 30/0 and 60/0). Closed bars: PPDE>0.95 (significant data). Open bars: PPDE<0.95 (less significant). Plots are derived from data in “Table S1 Transcriptome Data.xlsx”. HGNC Gene codes are displayed above the relative transcript level bars.

**Table 3.** GSEA parameters for Centromere Structure and Function, comparing stressed undifferentiated HL-60/S4 cells to unstressed control cells: NES, Normalized Enrichment Score; Nominal p, statistical significance, where p = 0.00 is the maximal significance and p > 0.05 is regarded as less significant.

| Gene Set | 30 min vs 0 min |  | 60 min vs 0 min |  | # of Genes |
| --- | --- | --- | --- | --- | --- |
|  | NES | Nominal p | NES | Nominal p |  |
| REACTOME_DEPOSITION_OF_NEW_CENPA_CONTAINING_NUCLEOSOMES_AT_THE_CENTROMERE | 2.02 | 0.0 | 2.08 | 0.0 | 69 |
| GOCC_CHROMOSOME_CENTROMERIC_CORE_DOMAIN | 1.49 | 0.03 | 1.70 | 0.01 | 18 |
| GOBP_CENTROMERE_COMPLEX_ASSEMBLY | 1.28 | 0.11 | 1.29 | 0.09 | 30 |
| GOCC_CHROMOSOME_CENTROMERIC_REGION | 1.08 | 0.16 | 1.09 | 0.15 | 259 |

### Hyperosmotic Stress with Sucrose maintains the individuality and integrity of the Chromosome Territories

In the interphase nucleus, it has been convincingly demonstrated, by employing “FISH” (Fluorescent In Situ Hybridization), that individual chromosomes are <u>not</u> dispersed around the entire nucleus, but are localized into distinct individual nuclear domains, called “Chromosome Territories” (Cremer and Cremer, 2010). Figure 8 presents the results of FISH experiments on undifferentiated HL-60/S4 cells. It is clear that for three different chromosomes (chr. 4,12 and 19) their distinct individuality is maintained within interphase nuclei and mitotic chromosomes, <u>even</u> following the dehydration shrinkage effect of hyperosmotic 300 mM sucrose. Individual chromosome territories do appear to have shrunken somewhat, in response to the osmotic stress. But a firm conclusion must await a more systematic study.

**Figure 8.**
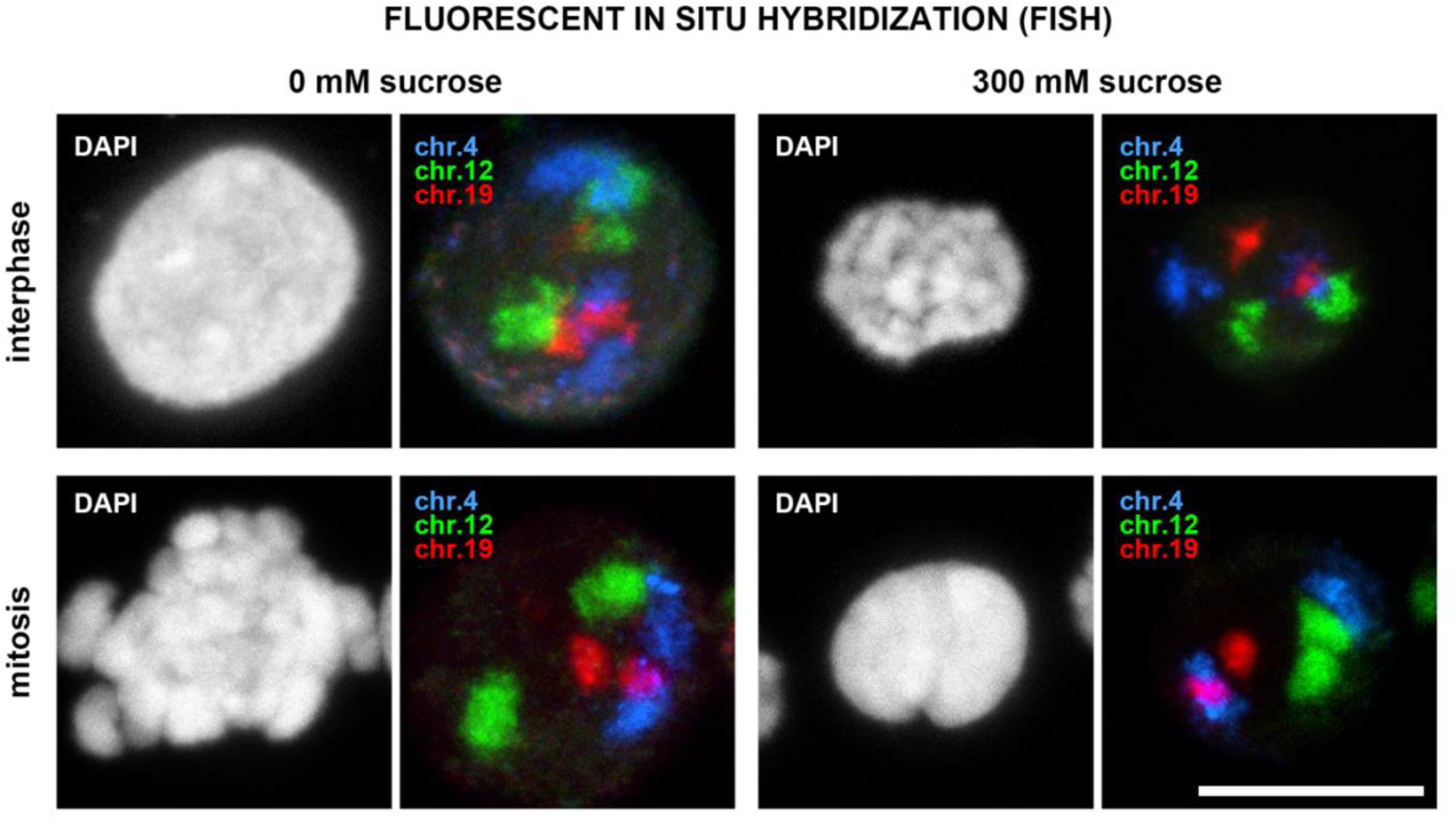
FISH localization of chromosome territories for chr. 4, 12 and 19 fixed in undifferentiated HL-60/S4 cells, incubated in iso-osmotic or hyperosmotic tissue culture medium. Magnification bar: 5 μm

### Hyperosmotic Stress with Sucrose produces increased Ribosome Biogenesis and mRNA Translation

Given the extensive amount of interphase chromatin congelation resulting from hyperosmotic stress (Figure 2), we had expected that many cellular functions would experience an almost immediate cessation. But, to our surprise, eight of the “Top 10” GSEA gene sets (i.e., exhibiting the highest NES values and p-scores of 0.0) are related to increased ribosome biosynthesis and function. Figure 9 presents four of these eight GSEA enrichment plots, each comparing the phenotypes at 30 *versus* 0 minutes of sucrose exposure. Table 4 summarizes the eight GSEA parameters for 30 *versus* 0 minutes, and for 60 *versus* 0 minutes. The implication of these results is that the nucleolus remains “active” during this time frame, assembling nascent ribosomes; i.e., the shrunken cells are continuing to build protein “structures” for, at least, 60 minutes. Evidently, this is a subset of the total mRNAs that exhibit <u>increased</u> synthesis, the opposite of mRNAs described earlier (Figure 3 and Table 1).

**Figure 9.**
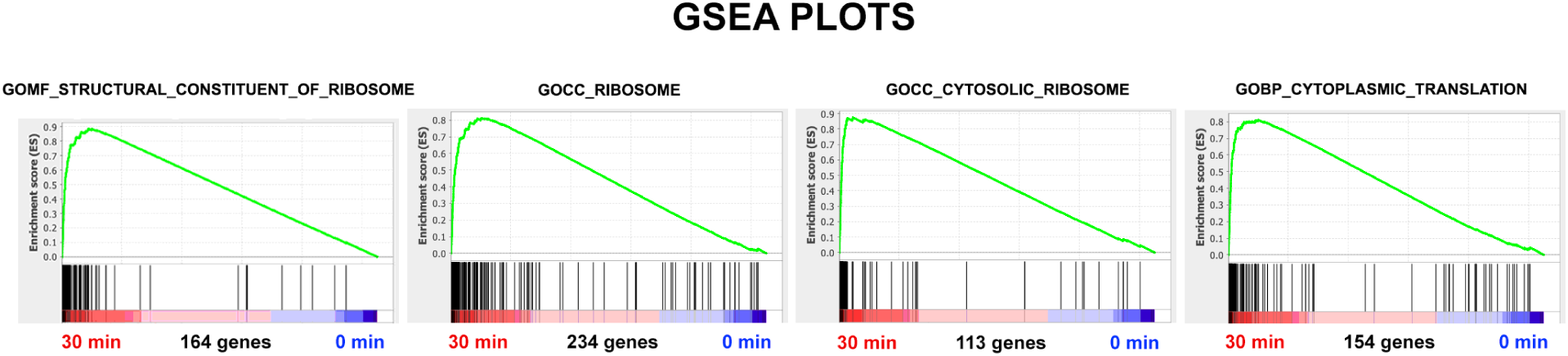
GSEA enrichment plots of Ribosome Biogenesis related transcripts demonstrating enrichment after 30 minutes of exposure to Hyperosmotic stress conditions, comparing stressed (30 min) undifferentiated HL-60/S4 cells to unstressed (0 min) control cells.

### The Nucleolus is a functioning structure during Hyperosmotic Stress with Sucrose

Figure 10 presents images of interphase HL-60/S4 cells, before and after 30 minutes of exposure to 300 mM sucrose, with emphasis upon nucleolar morphology. The top row of panels demonstrates that NPM1 (a ribosomal protein chaperon; www.genecards.org) outlines the nucleolar surface (+/-) sucrose treatment. PL2-6 continues to immunostain the surface of the amorphous chromosomes and the surface of chromatin at the nuclear envelopes. It has been shown that PL2-6 reacts with “exposed” nucleosomal “acidic patches” (Olins and Olins, 2018; Zhou et al., 2019). The bottom row of panels demonstrates that nucleoli (+/- sucrose treatment) are strongly stained for RNA, employing Syto RNASelect (ThermoFisher Scientific).

**Figure 10.**
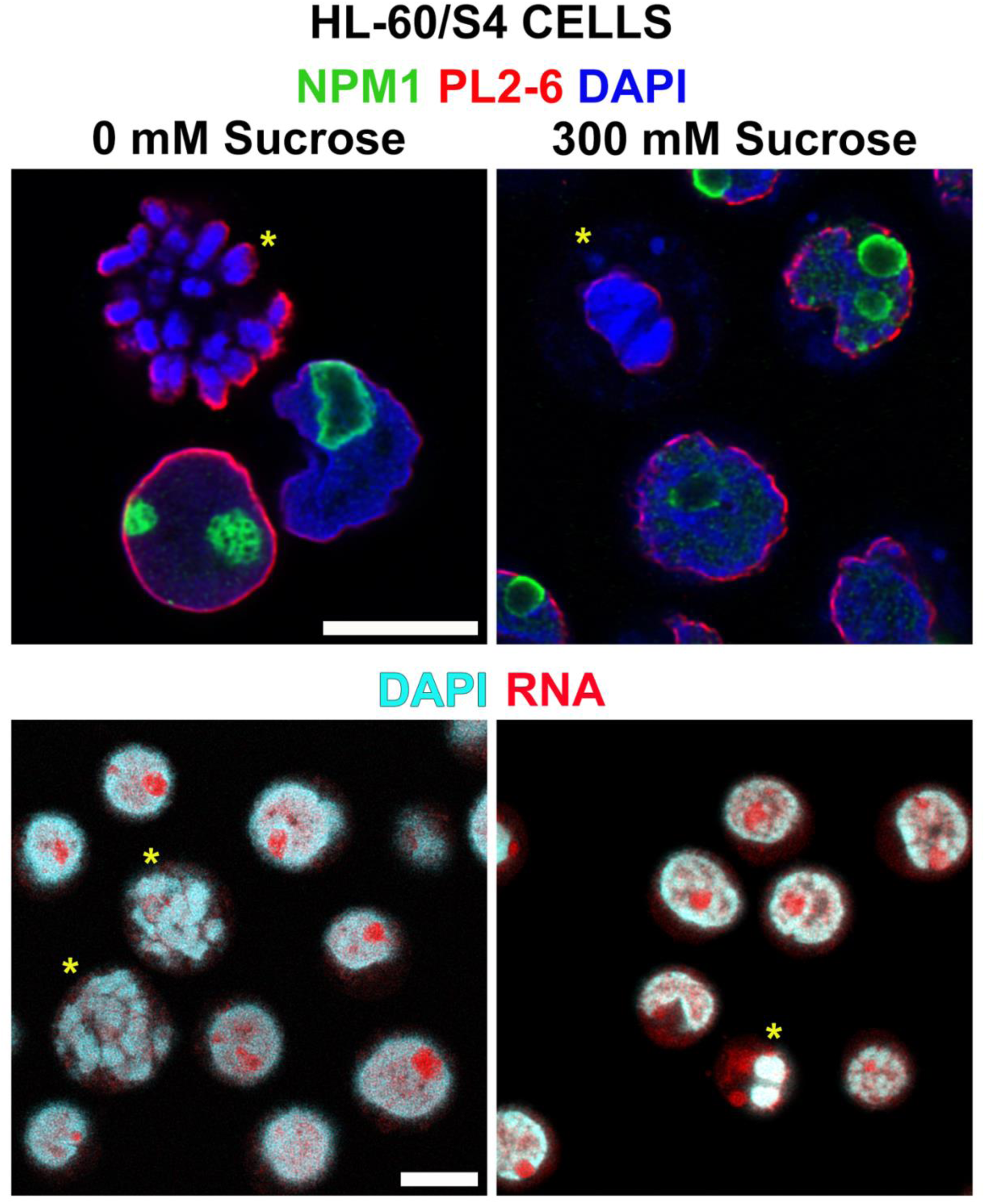
presents images of interphase HL-60/S4 cells, before and after 30 minutes of exposure to 300 mM sucrose. The yellow asterisks denote the positions of mitotic cells before and after exposure to hyperosmotic stress. Top row: Immunostaining of nucleoli (NPM1), epichromatin (PL2-6) and DNA (DAPI). Bottom row: Nuclear RNA (especially nucleoli) stained with Syto RNASelect. Magnification bars: 10 μm

### Hyperosmotic Stress with Sucrose produces increased transcripts for Mitochondrial Oxidative Phosphorylation and ATP biosynthesis

Ten of the “Top 20” GSEA gene sets are related to mitochondrial structure and function. The implication of their enrichment with mitochondrial protein transcripts is that ATP biosynthesis is very likely to be increased during cell dehydration (Figure 11 and Table 5). Figure 12 confirms that the mitochondria are “active” in both 0- and 300-mM sucrose.

**Table 4.** GSEA parameters for Ribosome Biogenesis, comparing stressed undifferentiated HL-60/S4 cells to unstressed control cells: NES, Normalized Enrichment Score; Nominal p, statistical significance, where Nominal p = 0.00 is the maximal significance and Nominal p > 0.05 is regarded as less significant.

| Gene Set | 30 min vs 0 min |  | 60 min vs 0 min |  | # of Genes |
| --- | --- | --- | --- | --- | --- |
|  | NES | Nominal p | NES | Nominal p |  |
| **GOCC_RIBOSOMAL_SUBUNIT** |  |  |  |  |  |
|  | 2.87 | 0.0 | 2.78 | 0.0 | 183 |
| **GOMF_STRUCTURAL_CONSTITUENT_OF_RIBOSOME** |  |  |  |  |  |
|  | 2.86 | 0.0 | 2.84 | 0.0 | 164 |
| **GOCC_RIBOSOME** |  |  |  |  |  |
|  | 2.73 | 0.0 | 2.67 | 0.0 | 234 |
| **GOCC_CYTOSOLIC_RIBOSOME** |  |  |  |  |  |
|  | 2.72 | 0.0 | 2.64 | 0.0 | 113 |
| **GOCC_LARGE_RIBOSOMAL_SUBUNIT** |  |  |  |  |  |
|  | 2.72 | 0.0 | 2.67 | 0.0 | 114 |
| **GOBP_CYTOPLASMIC_TRANSLATION** |  |  |  |  |  |
|  | 2.59 | 0.0 | 2.56 | 0.0 | 154 |
| **GOCC_ORGANELLAR_RIBOSOME** |  |  |  |  |  |
|  | 2.57 | 0.0 | 2.54 | 0.0 | 89 |
| **GOCC_SMALL_RIBOSOMAL_SUBUNIT** |  |  |  |  |  |
|  | 2.53 | 0.0 | 2.52 | 0.0 | 73 |

**Table 5.** GSEA parameters for Mitochondrial Function, comparing stressed undifferentiated HL-60/S4 cells to unstressed control cells: NES, Normalized Enrichment Score; Nominal p, statistical significance, where p = 0.00 is the maximal significance and p > 0.05 is regarded as less significant.

| Gene Set | 30 min vs 0 min |  | 60 min vs 0 min |  | # of Genes |
| --- | --- | --- | --- | --- | --- |
|  | NES | Nominal p | NES | Nominal p |  |
| **GOCC_MITOCHONDRIAL_PROTEIN_CONTAINING_COMPLEX** |  |  |  |  |  |
|  | 2.72 | 0.0 | 2.68 | 0.0 | 280 |
| **GOBP_ATP_SYNTHESIS_COUPLED_ELECTRON_TRANSPORT** |  |  |  |  |  |
|  | 2.56 | 0.0 | 2.44 | 0.0 | 87 |
| **GOCC_INNER_MITOCHONDRIAL_MEMBRANE_PROTEIN_COMPLEX** |  |  |  |  |  |
|  | 2.55 | 0.0 | 2.53 | 0.0 | 138 |
| **GOBP_MITOCHONDRIAL_TRANSLATION** |  |  |  |  |  |
|  | 2.55 | 0.0 | 2.51 | 0.0 | 130 |
| **GOCC_RESPIRASOME** |  |  |  |  |  |
|  | 2.53 | 0.0 | 2.45 | 0.0 | 87 |
| **GOBP_MITOCHONDRIAL_RESPIRATORY_CHAIN_COMPLEX_ASSEMBLY** |  |  |  |  |  |
|  | 2.51 | 0.0 | 2.50 | 0.0 | 92 |
| **GOBP_OXIDATIVE_PHOSPHORYLATION** |  |  |  |  |  |
|  | 2.50 | 0.0 | 2.42 | 0.0 | 130 |
| **GOBP_AEROBIC_RESPIRATION** |  |  |  |  |  |
|  | 2.47 | 0.0 | 2.38 | 0.0 | 179 |
| **GOBP_MITOCHONDRIAL_GENE_EXPRESSION** |  |  |  |  |  |
|  | 2.44 | 0.0 | 2.37 | 0.0 | 163 |
| **GOCC_MITOCHONDRIAL_LARGE_RIBOSOMAL_SUBUNIT** |  |  |  |  |  |
|  | 2.39 | 0.0 | 2.39 | 0.0 | 56 |

**Figure 11.**
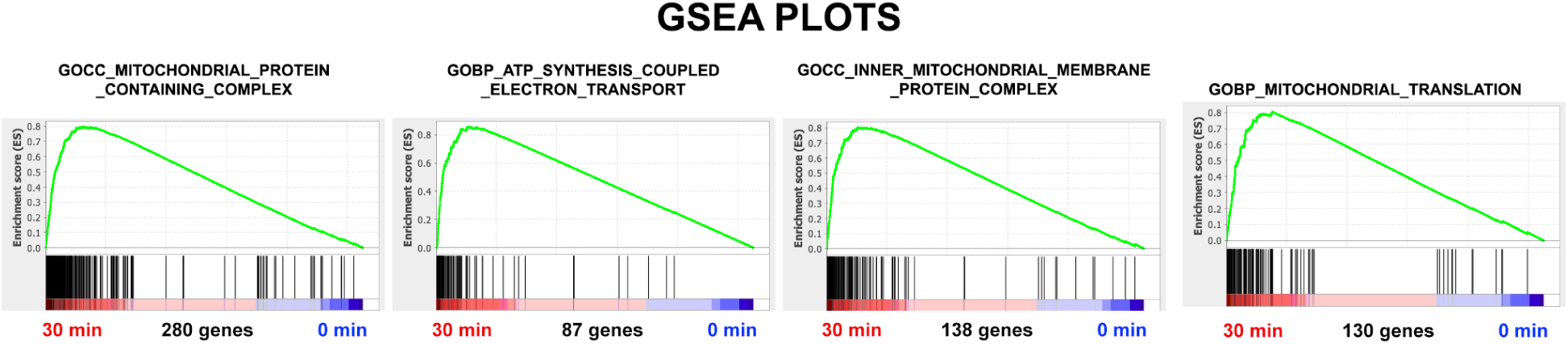
GSEA enrichment plots of Mitochondrial Function related transcripts demonstrating enrichment after 30 minutes of exposure to Hyperosmotic stress conditions, comparing stressed undifferentiated HL-60/S4 cells to unstressed (0 min) control cells.

**Figure 12.**
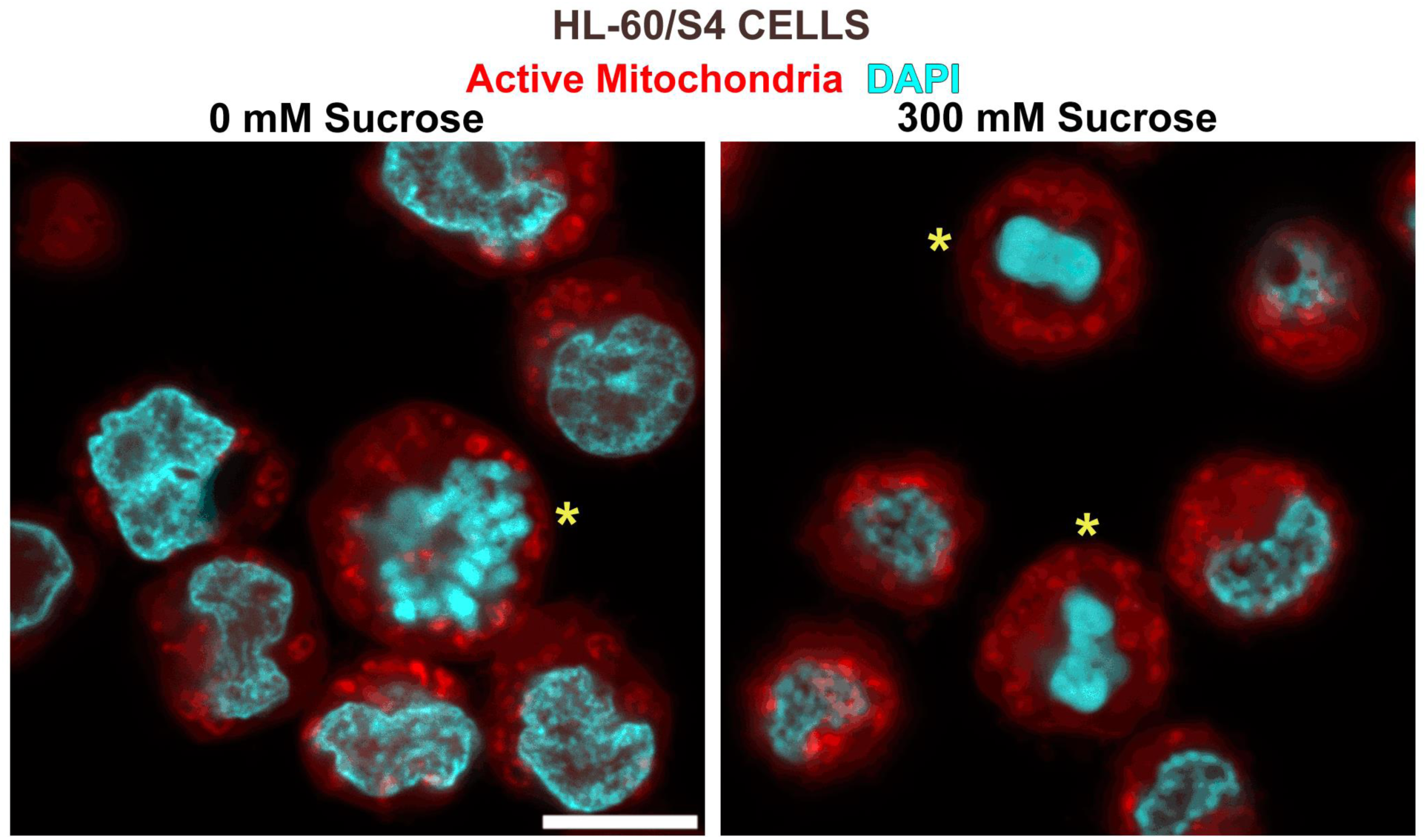
*In Vivo* staining by MitoTracker Red of undifferentiated HL-60/S4 mitochondria in 0 and 300 mM sucrose for 30 min, prior to fixation with HCHO. The yellow asterisks denote the positions of mitotic cells before and after exposure to hyperosmotic stress. Colors: Active Mitochondria (Red); DNA (Cyan). The red staining of mitochondria indicates that they were active in both buffer situations. Magnification bar: 10 μm

### Hyperosmotic Stress with Sucrose produces enrichment of Proteasome gene transcripts supporting increased protein turnover

**Figure 13.**
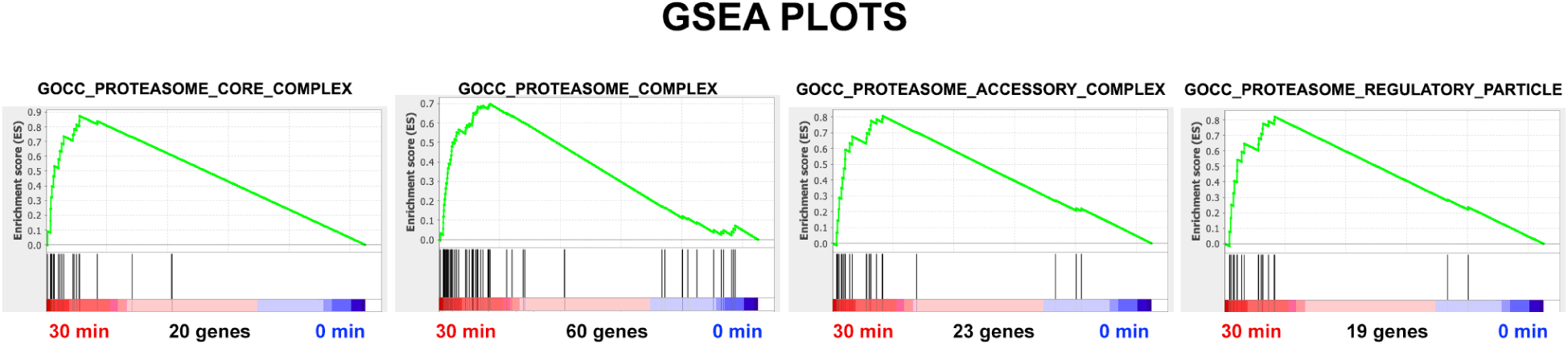
GSEA enrichment plots of Proteasome Activity related transcripts demonstrating enrichment after 30 minutes of exposure to Hyperosmotic stress conditions, comparing stressed undifferentiated HL-60/S4 cells to unstressed (0 min) control cells.

**Table 6.** GSEA parameters for Proteasome Activity, comparing stressed undifferentiated HL-60/S4 cells to unstressed control cells: NES, Normalized Enrichment Score; Nominal p, statistical significance, where p = 0.00 is the maximal significance and p > 0.05 is regarded as less significant.

| Gene Set | 30 min vs 0 min |  | 60 min vs 0 min |  | # of Genes |
| --- | --- | --- | --- | --- | --- |
|  | NES | Nominal p | NES | Nominal p |  |
| **GOCC_PROTEASOME_CORE_COMPLEX** |  |  |  |  |  |
|  | 2.00 | 0.0 | 1.89 | 0.0 | 20 |
| **GOCC_PROTEASOME_COMPLEX** |  |  |  |  |  |
|  | 2.00 | 0.0 | 1.86 | 0.0 | 60 |
| **GOCC_PROTEASOME_ACCESSORY_COMPLEX** |  |  |  |  |  |
|  | 1.89 | 0.0 | 1.78 | 0.0 | 23 |
| **GOCC_PROTEASOME_REGULATORY_PARTICLE** |  |  |  |  |  |
|  | 1.87 | 0.0 | 1.71 | 0.0 | 19 |

### Hyperosmotic Stress with Sucrose produces relocation of LBR from the nuclear envelope to the cytoplasm

Lamin B Receptor (LBR) normally resides in the interphase nuclear envelope, tethered to the inner nuclear membrane via a series of C-terminal transmembrane segments (amino acid residues ∼200-615) (Olins et al., 2010b). LBR has an N-terminal (Tudor Domain) peptide segment (1-60 amino acids) proximal to a peptide region (∼97-124 aa) bound to Heterochromatin Protein 1 α (CBX5) (Olins et al., 2010b). The Tudor Domain binds to the histone H4K20me2 epigenetic marker (Hirano et al., 2012; Olins et al., 2010b). The CBX5 “Chromo Domain” binds to histone H3K9me2 and me3 heterochromatin epigenetic markers (Strom et al., 2021). The CBX5 “Chromo Shadow Domain” binds to LBR residues 97-174) (Olins et al., 2010b). Thus, LBR forms bridges from the nuclear envelope to the underlying heterochromatin.

When undifferentiated HL-60/S4 cells are exposed to hyperosmotic stress with 300 mM sucrose in tissue culture medium, besides the cell shrinkage and chromatin congelation, LBR exits the nuclear envelope and spreads throughout the cytoplasm (Figure 14 top panels). Presumably, the interactions between the LBR N-termini and the heterochromatin epigenetic marks have been severed, liberating LBR to diffuse broadly in membranous space. A similar dispersion of CBX5 into the cytoplasm after hyperosmotic stress is also observed (Figure 14 bottom panels), but CBX5 does <u>not</u> possess transmembrane segments, like LBR. At present, it is unknown whether CBX5 is co-localized and interacting with LBR, when both are present within the cytoplasm.

**Figure 14.**
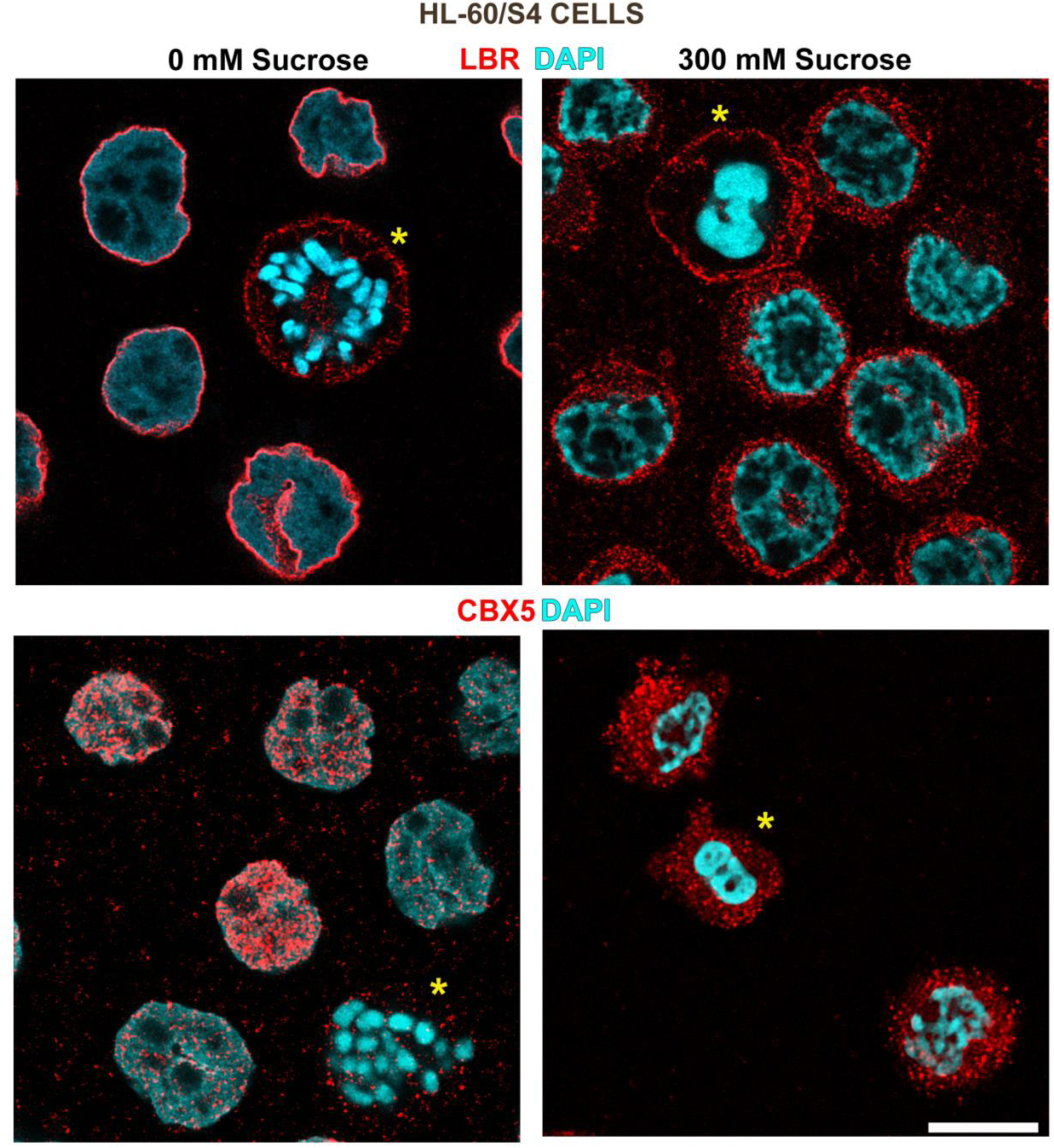
Hyperosmotic Stress of undifferentiated HL-60/S4 cells: Sucrose induced relocation and dispersion into the cytoplasm of LBR (top panels) and of CBX5 (bottom panels). Colors: LBR and CBX5 (red); DNA (cyan). The yellow asterisks denote the positions of mitotic cells before and after exposure to hyperosmotic stress. CBX5 is affiliated with interphase chromatin, but not with mitotic chromosomes of unstressed cells. Magnification bar: 10 μm

HL-60/S4 cells can be differentiated into granulocytes by treatment with retinoic acid (RA). The granulocytes have multilobed nuclei connected by chromatin sheets (ELCS or Envelope-Limited Chromatin Sheets) (Eltsov et al., 2014; Olins et al., 1998; Xu et al., 2021). These changes in nuclear shape depend upon an increase of cellular LBR. LBR knockdown prevents the RA-induced nuclear shape changes (i.e., nuclear lobulation and ELCS formation) (Mark Welch et al., 2024; Olins et al., 2010a).

When HL-60/S4 granulocytes are exposed to hyperosmotic stress with sucrose, the cells shrink and LBR migrates into the cytoplasm, as observed with undifferentiated HL-60/S4 cells (Figure 15). However, some nuclear lobulation and ELCS are maintained in the 300 mM sucrose, when immunostained with anti-LMNB2 (Figure 16). This apparent stability may occur in the earlier expansion of nuclear envelope membranes during RA-induced granulocytic differentiation (Olins et al., 1998). Stabilization might result from the nuclear envelope membranes binding to lamina (e.g., LMNB2) and/or higher-order chromatin structure (e.g., 30 nm fibers) associated with the nuclear envelope (Eltsov et al., 2014; Olins and Olins, 2009; Xu et al., 2021).

**Figure 15.**
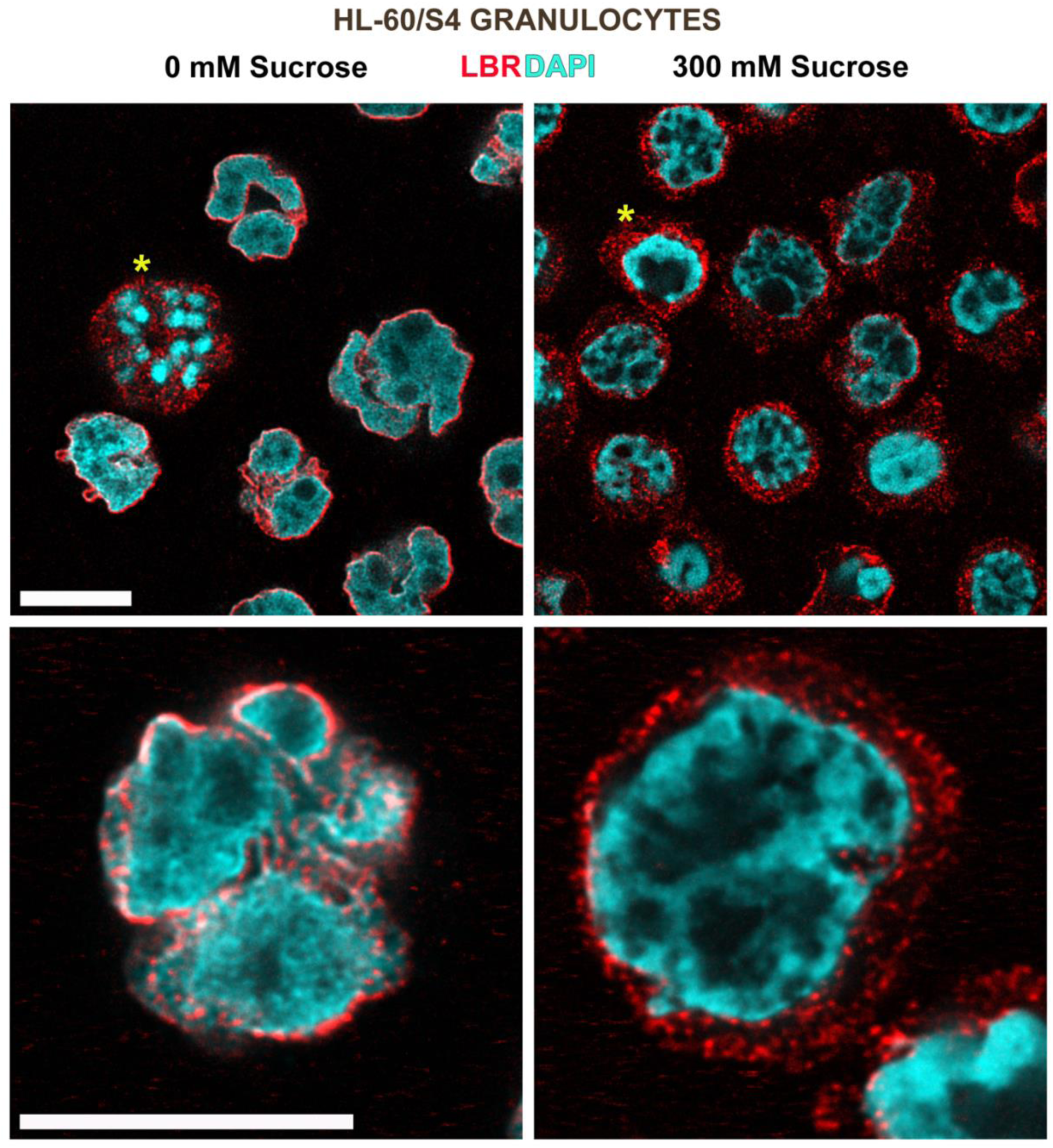
Hyperosmotic Stress (300 mM sucrose) on RA-treated granulocytic HL-60/S4 cell induced dispersion of LBR into the cytoplasm. ELCS (stained with anti-LBR) can be visualized spanning between nuclear lobes in 0 mM sucrose. Colors: LBR (red); DNA (cyan). The yellow asterisks denote the positions of mitotic cells before and after exposure to hyperosmotic stress. Magnification bars: 10 μm

**Figure 16.**
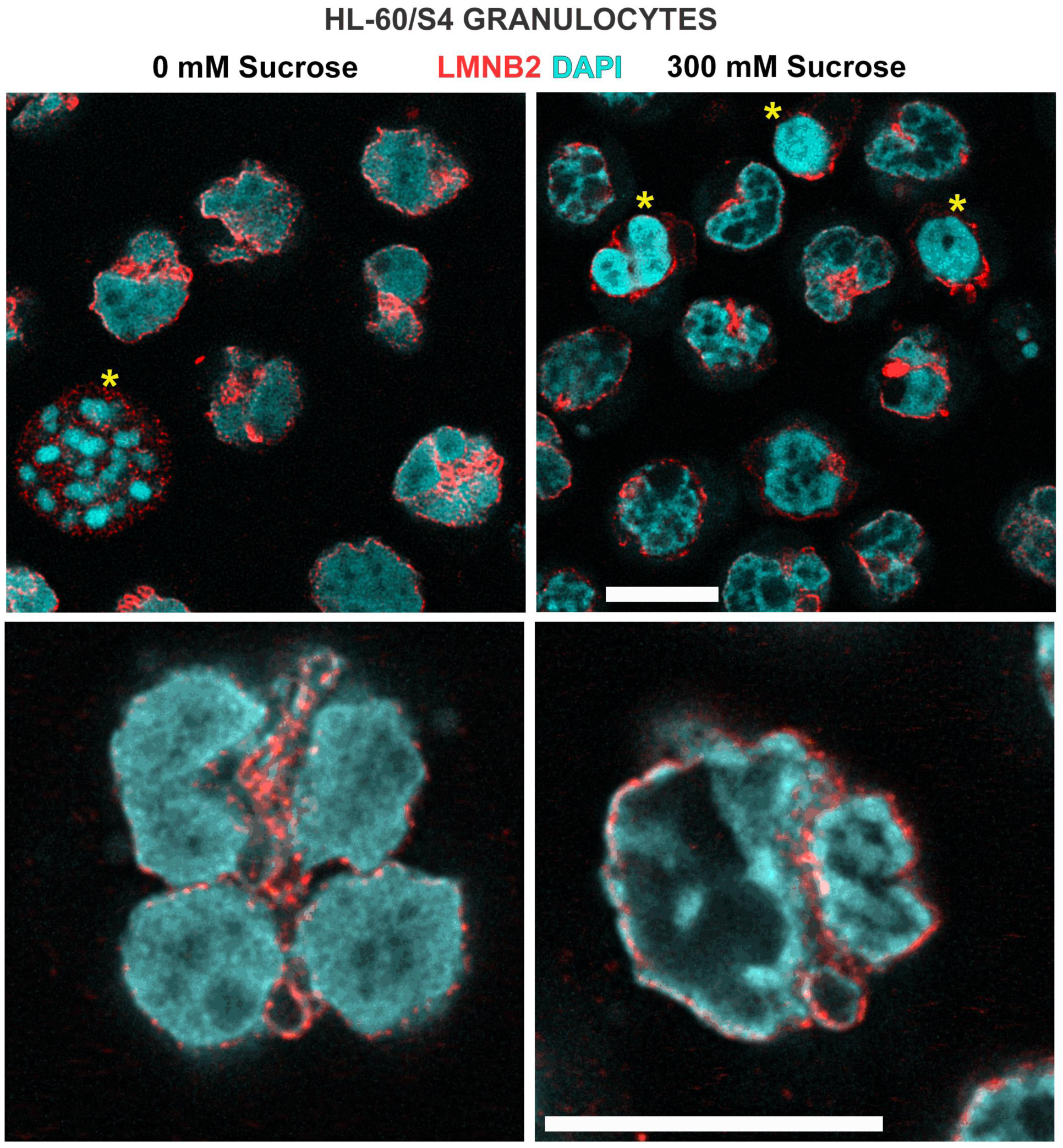
Hyperosmotic Stress (300 mM sucrose) of RA-treated granulocytic HL-60/S4 cells does <u>not</u> induce dispersion of Lamin B2 (LMNB2) into the cytoplasm. ELCS continue to retain LMNB2. Some ELCS appear to be congealed in 300 mM sucrose. Colors: LMNB2 (red); DNA (cyan). The yellow asterisks denote the positions of mitotic chromosomes before and after exposure to hyperosmotic stress. Magnification bars: 10 μm

### Hyperosmotic Stress downregulates transcripts for DNA methylation and for methylated DNA binding proteins

DNA methylation at CpG sites is associated with gene repression and heterochromatin formation (Ghosh et al., 2010; Hansen et al., 2010; Lee et al., 2020; Ortega-Alarcon et al., 2024). The effect of hyperosmotic stress is dramatically demonstrated by examining gene sets. (Left): GOBP_DNA_METHYLATION_DEPENDENT_HETEROCHROMATIN_FORMATION The Leading Edge of this gene set reveals the downregulation of two important genes: DNMT1 and DNMT3A (DNA methyltransferases), enzymes critical for DNA methylation. (Middle and Right): REACTOME_REGULATION_OF_MECP2_EXPRESSION_AND_ACTIVITY and REACTOME_TRANSCRIPTIONAL_REGULATION_BY_MECP2. Both gene sets exhibit downregulation of LBR, MECP2, NCOR1 and NCOR2. The latter two genes are nuclear receptor co-repressors and ligands of MECP2. From these few examples of downregulated gene expression during hyperosmotic stress, we hypothesize that DNA methylation is compromised, adversely affecting the binding of repressor and heterochromatin-associated proteins. This apparent reduction of DNA methylation by hyperosmotic stress correlates with the decrease in heterochromatin described earlier (Figure 3 and Table 1).

**Figure 17.**
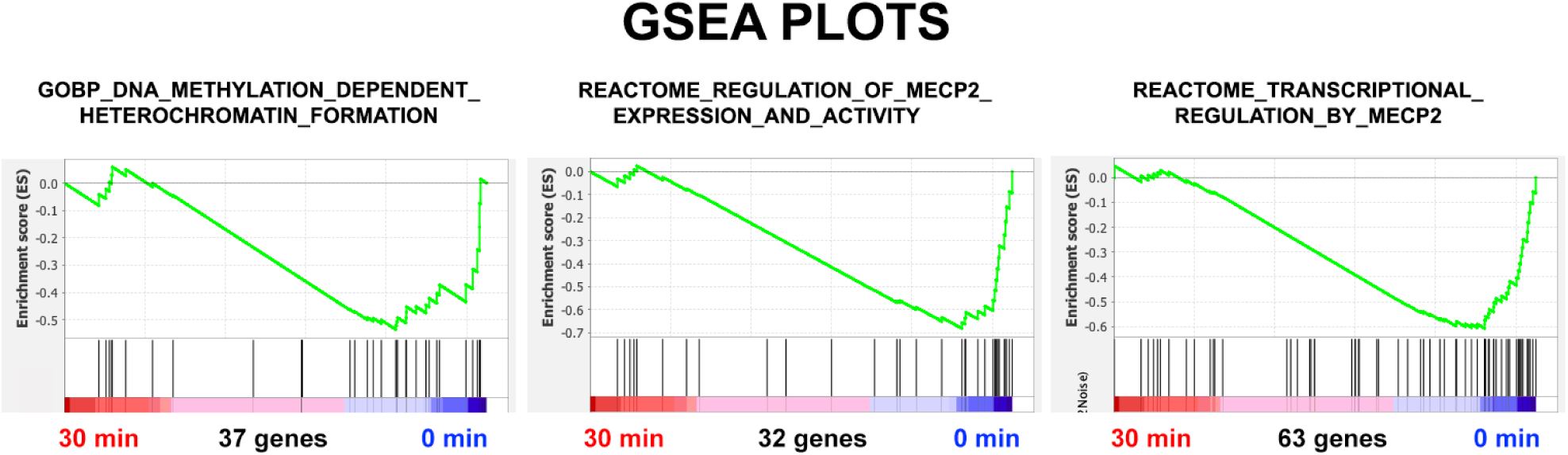
GSEA enrichment plots of genes relevant to DNA methylation and to gene regulation by MECP2, following 30 minutes of Hyperosmotic stress conditions (300 mM sucrose), comparing stressed undifferentiated cells to unstressed HL-60/S4 control (0) cells.

**Table 7.** GSEA parameters for DNA methylation and MECP activity, comparing stressed undifferentiated HL-60/S4 cells to unstressed control cells: NES, Normalized Enrichment Score; Nominal p, statistical significance, where p = 0.00 is the maximal significance and p > 0.05 is regarded as less significant.

| Gene Set | 30 min vs 0 min |  | 60 min vs 0 min |  | # of Genes |
| --- | --- | --- | --- | --- | --- |
|  | NES | Nominal p | NES | Nominal p |  |
| GOBP_DNA_METHYLATION_DEPENDENT_HETEROCHROMATIN_FORMATION | -1.31 | 0.10 | -1.51 | 0.02 | 37 |
| REACTOME_REGULATION_OF_MECP2_EXPRESSION_AND_ACTIVITY | -1.63 | 0.01 | -1.61 | 0.01 | 32 |
| REACTOME_TRANSCRIPTIONAL_REGULATION_BY_MECP2 | -1.62 | 0.0 | -1.61 | 0.0 | 63 |

With regard to chromatin structure, MECP2 is a particularly interesting protein. MECP2 binds to methylated DNA sites, employing the “methyl binding domain” (MBD). In addition, MECP2 binds to histones and competes with histone H1 binding to the nucleosome. MECP2 is an Intrinsically Disordered Protein, with the capabilities of promiscuous interactions (Ortega-Alarcon et al., 2024). Immunostaining MeCP2 in undifferentiated and granulocytic HL-60/S4, with-or-without hyperosmotic stress from sucrose (30 minutes) are presented in Figures 18 and 19. In 0 mM sucrose, MECP2 appears dispersed within the interphase chromatin, possibly due to the exposure of many methylated DNA binding sites. In 300 mM sucrose, MECP2 seems to be segregated from congealed chromatin, perhaps due to a reduction of methylated DNA binding sites.

**Figure 18.**
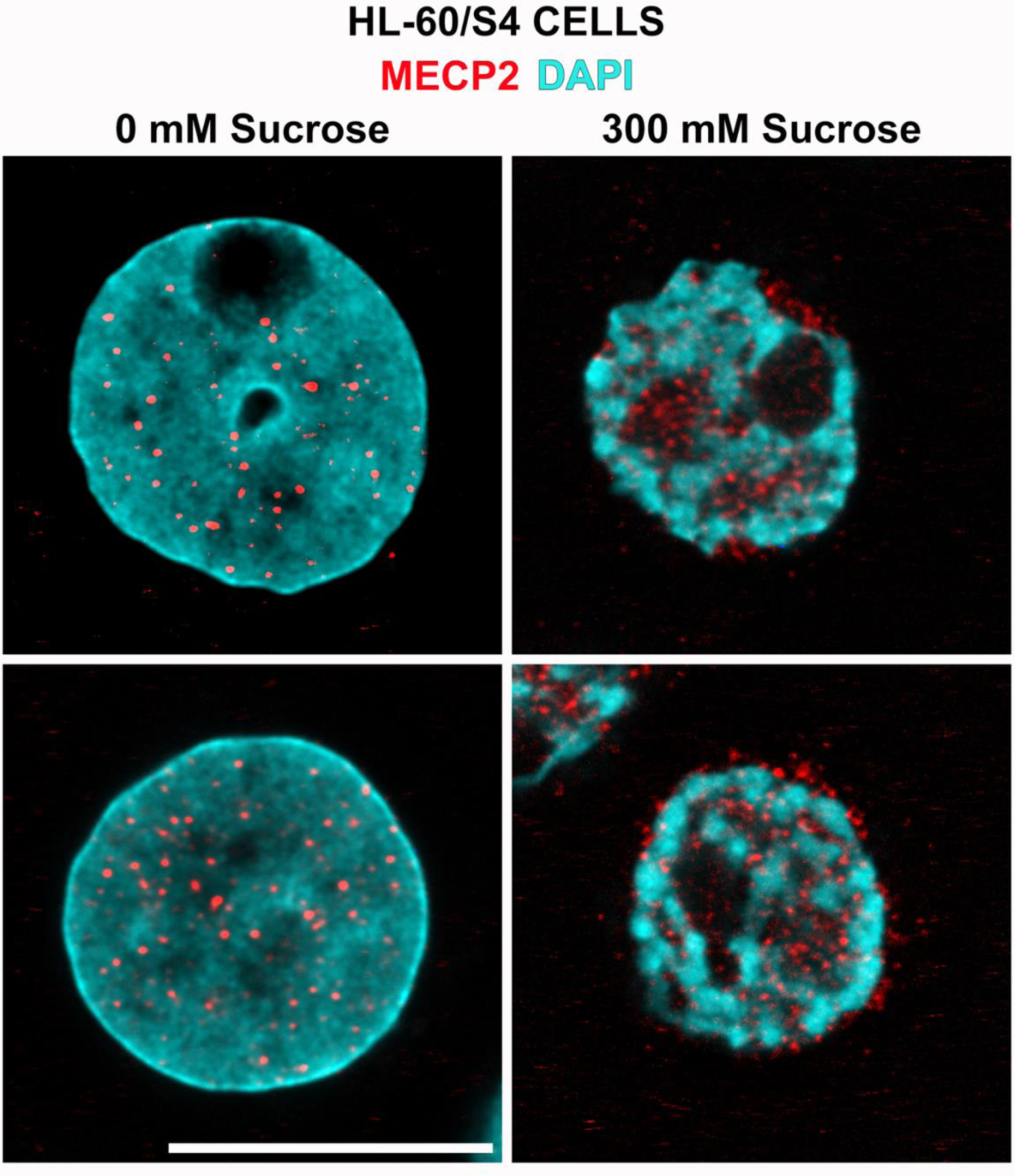
Hyperosmotic Stress of undifferentiated HL-60/S4 cells stained with anti-MECP2 (red) and DNA (cyan). MECP2 appears dispersed within the chromatin of unstressed interphase nuclei (left column, 0 mM sucrose); but appears segregated at the surfaces and within “voids” of congealed interphase chromatin (right column, 300 mM sucrose). Magnification bars: 10 μm

**Figure 19.**
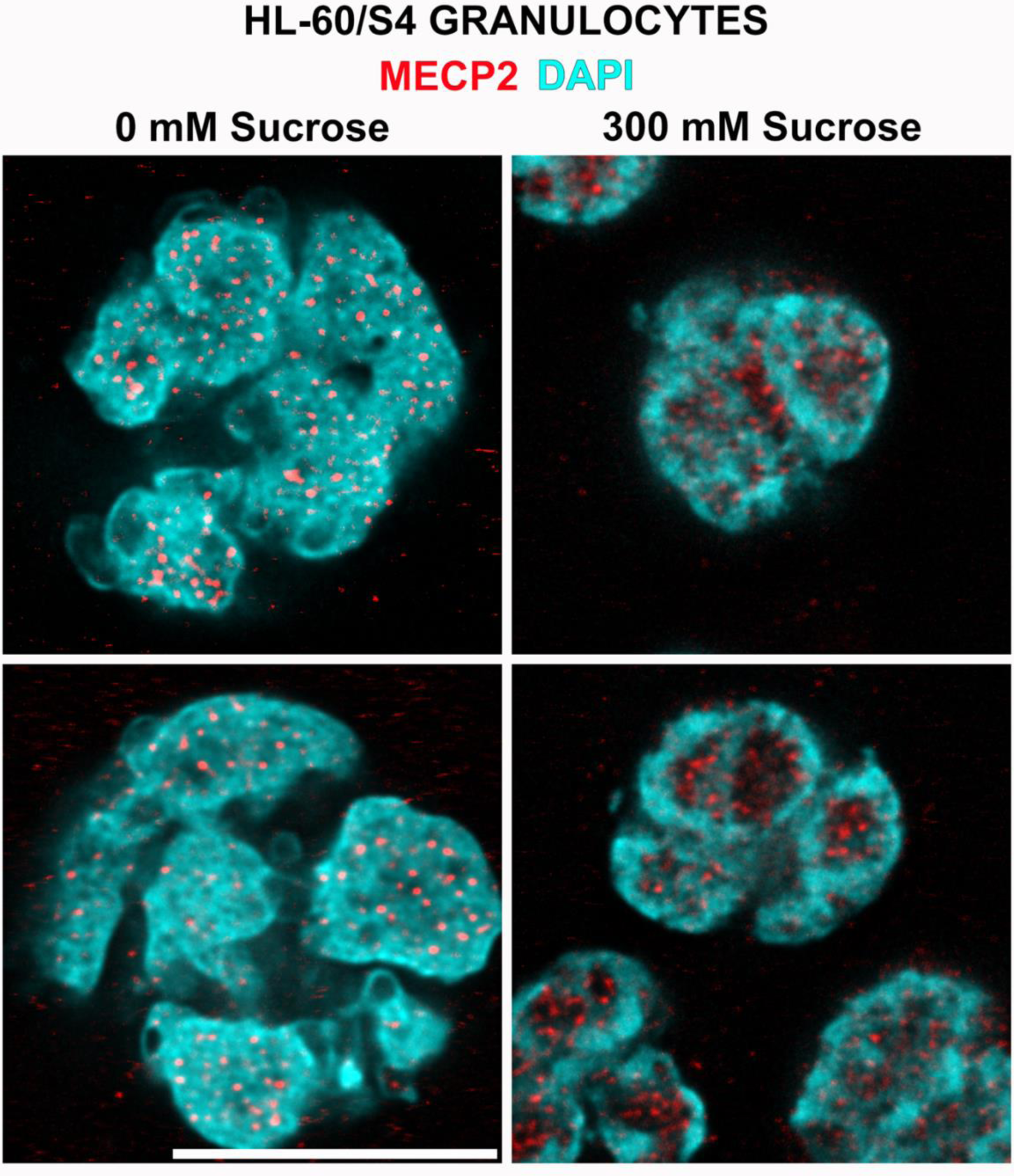
Hyperosmotic Stress of RA-treated granulocytic HL-60/S4 cells stained with anti-MECP2 (red) and DNA (cyan). MECP2 appears dispersed within the chromatin of unstressed nuclear lobules (left column, 0 mM sucrose); but appears segregated into “voids” of congealed lobular chromatin (right column, 300 mM sucrose). Magnification bars: 10 μm

## Discussion

The present article is our third, following (Olins et al., 2020) and (Mark Welch et al., 2022), principally concerned with the effects of acute hyperosmotic sucrose stress upon undifferentiated HL-60/S4 myeloid cells. The first article (Olins et al., 2020) was primarily an immunostaining microscopic study, which demonstrated the congelation of interphase chromatin and mitotic chromosomes, frequently accompanied by phase separation of various chromatin components.

The second article (Mark Welch et al., 2022) documented the changes in gene transcription, comparing 30 and 60 minutes of hyperosmotic sucrose stress to the untreated undifferentiated control HL-60/S4 cells. Functional analysis of the transcripts (in the second article) employed overabundance analysis of Gene Ontology (GO) gene sets. The present article employs the same transcript abundance data, but uses Gene Set Enrichment Analysis (GSEA) (Mootha et al., 2003; Subramanian et al., 2005) to predict the enrichment of phenotypes. In addition, the present immunostaining microscopy identifies the localization of key proteins in nuclei of different phenotypes. Furthermore, the present article examines the microscopic consequences of hyperosmotic sucrose stress on HL-60/S4 RA-induced granulocytes (Figures 15, 16 and 19). Similar stress-induced relocation of specific epitopes was observed in both undifferentiated and granulocytic forms of HL-60/S4 cells.

Comparing the transcript overabundance and the GSEA results, there is general agreement that heterochromatin and histone lysine methyltransferase exhibit reduction of their related transcripts. There is also agreement that genes associated with ribosome biosynthesis, mitochondrial and proteosome activity have increased transcript levels. In our earlier article (Mark Welch et al., 2022), we observed that Replication-Dependent Histone mRNAs are over-represented at 30 and 60-minutes of sucrose stress, even though these cells have ceased dividing. Combining these earlier observations with the present article, we are struck by the <u>lack</u> of “meaningful phenotypic coordination”, a property characteristic of HL-60/S4 cells induced to differentiate to granulocytes with retinoic acid (RA) or to macrophages with phorbol ester (TPA) (Mark Welch et al., 2017). Compared to HL-60/S4 cell differentiation, the hyperosmotic stress response appears to be chaotic in the phenotypic changes. Quoting from our earlier article: “The induced physiological state of acute hyperosmotically stressed undifferentiated HL-60/S4 cells resembles a rapid attempt to rebuild the damaged cells …ultimately unable to prevent cell death.” (Mark Welch et al., 2022).

The biophysical changes that occur in hyperosmotic stressed cells with consequent changes in nuclear and chromatin interactions are of considerable interest (Hancock, 2014). There are numerous studies on the biophysical consequences of osmotic stress; e.g., see (Kitamura et al., 2023; Thiemicke and Neuert, 2021; Watanabe et al., 2021). HL-60/S4 cells apparently do not synthesize sufficient osmolytes to counter the imposed osmotic pressure from 300 mM sucrose, as evidenced by the cell shrinkage and chromatin congelation with apparent *in situ* phase separation (Olins et al., 2020). *In situ* chromatin is a very heterogeneous structure; “walking along” a chromosome, one encounters a changing composition of nonhistones and epigenetic markings. It is to be expected that these contiguous heterogeneous chromatin micro-environments will respond very differently to hyperosmotic stress, compared to regulation by normal programmed cell differentiation factors. Furthermore, hyperosmotic stress may deliver abnormal and/or poorly coordinated signals to gene regulatory chromatin regions. If this perspective is correct, it is reasonable to suggest that, as a consequence of hyperosmotic stress, control of genetic expression can be profoundly disabled with consequent disruption of normal cellular homeostasis.

## Supplemental Files

**Table S1** Transcriptome Data

**Table S2** Formatted for Expression dataset

**Table S3** Methyltransferase Leading Edges

## Acknowledgements

The authors wish to express our gratitude to Dr. Marion Cremer and Dr. Irina Solovei, Biozentrum, LMU, 82152 Martinsried, Germany, for conducting the FISH experiments (Figure 8). This project utilized the services of the Histopathology and Microscopy Core at the MaineHealth Institute for Research, which was supported by NIH/NIGMS P30GM106391 and P20GM121301 grants.

## References

Cremer, T., and M. Cremer. 2010. Chromosome territories. Cold Spring Harb Perspect Biol. 2:a003889.

Eltsov, M., S. Sosnovski, A.L. Olins, and D.E. Olins. 2014. ELCS in ice: cryo-electron microscopy of nuclear envelope-limited chromatin sheets. Chromosoma. 123:303–312.

Finan, J.D., and F. Guilak. 2010. The effects of osmotic stress on the structure and function of the cell nucleus. J Cell Biochem. 109:460–467.

Finan, J.D., H.A. Leddy, and F. Guilak. 2011. Osmotic stress alters chromatin condensation and nucleocytoplasmic transport. Biochem Biophys Res Commun. 408:230–235.

Ghosh, R.P., R.A. Horowitz-Scherer, T. Nikitina, L.S. Shlyakhtenko, and C.L. Woodcock. 2010. MeCP2 binds cooperatively to its substrate and competes with histone H1 for chromatin binding sites. Mol Cell Biol. 30:4656–4670.

Hancock, R. 2014. The crowded nucleus. Int Rev Cell Mol Biol. 307:15–26.

Hansen, J.C., R.P. Ghosh, and C.L. Woodcock. 2010. Binding of the Rett syndrome protein, MeCP2, to methylated and unmethylated DNA and chromatin. IUBMB Life. 62:732–738.

Hirano, Y., K. Hizume, H. Kimura, K. Takeyasu, T. Haraguchi, and Y. Hiraoka. 2012. Lamin B receptor recognizes specific modifications of histone H4 in heterochromatin formation. J Biol Chem. 287:42654–42663.

Irianto, J., J. Swift, R.P. Martins, G.D. McPhail, M.M. Knight, D.E. Discher, and D.A. Lee. 2013. Osmotic challenge drives rapid and reversible chromatin condensation in chondrocytes. Biophys J. 104:759–769.

Kitamura, A., S. Oasa, H. Kawaguchi, M. Osaka, V. Vukojević, and M. Kinjo. 2023. Increased intracellular crowding during hyperosmotic stress. Sci Rep. 13:11834.

Lee, W., J. Kim, J.M. Yun, T. Ohn, and Q. Gong. 2020. MeCP2 regulates gene expression through recognition of H3K27me3. Nat Commun. 11:3140.

Mark Welch, D.B., T.J. Gould, A.L. Olins, and D.E. Olins. 2022. The transcriptome of acute dehydration in Myeloid Leukemia cells. *bioRxiv*:2022.2009.2023.509183.

Mark Welch, D.B., A. Jauch, J. Langowski, A.L. Olins, and D.E. Olins. 2017. Transcriptomes reflect the phenotypes of undifferentiated, granulocyte and macrophage forms of HL-60/S4 cells. Nucleus. 8:222–237.

Mark Welch, D.B., A.L. Olins, and D.E. Olins. 2024. The Effects of Lamin B Receptor knockdown on a Myeloid Leukemia Cell. *bioRxiv*:2024.2006.2019.598074.

Mootha, V.K., C.M. Lindgren, K.F. Eriksson, A. Subramanian, S. Sihag, J. Lehar, P. Puigserver, E. Carlsson, M. Ridderstråle, E. Laurila, N. Houstis, M.J. Daly, N. Patterson, J.P. Mesirov, T.R. Golub, P. Tamayo, B. Spiegelman, E.S. Lander, J.N. Hirschhorn, D. Altshuler, and L.C. Groop. 2003. PGC-1alpha-responsive genes involved in oxidative phosphorylation are coordinately downregulated in human diabetes. Nat Genet. 34:267–273.

Olins, A.L., B. Buendia, H. Herrmann, P. Lichter, and D.E. Olins. 1998. Retinoic acid induction of nuclear envelope-limited chromatin sheets in HL-60. Exp Cell Res. 245:91–104.

Olins, A.L., A. Ernst, M. Zwerger, H. Herrmann, and D.E. Olins. 2010a. An in vitro model for Pelger-Huët anomaly: stable knockdown of lamin B receptor in HL-60 cells. Nucleus. 1:506–512.

Olins, A.L., T.J. Gould, L. Boyd, B. Sarg, and D.E. Olins. 2020. Hyperosmotic stress: in situ chromatin phase separation. Nucleus. 11:1–18.

Olins, A.L., G. Rhodes, D.B. Welch, M. Zwerger, and D.E. Olins. 2010b. Lamin B receptor: multi-tasking at the nuclear envelope. Nucleus. 1:53–70.

Olins, D.E., and A.L. Olins. 2009. Nuclear envelope-limited chromatin sheets (ELCS) and heterochromatin higher order structure. Chromosoma. 118:537–548.

Olins, D.E., and A.L. Olins. 2018. Epichromatin and chromomeres: a ‘fuzzy’ perspective. Open Biol. 8.

Ortega-Alarcon, D., R. Claveria-Gimeno, S. Vega, L. Kalani, O.C. Jorge-Torres, M. Esteller, J. Ausio, O. Abian, and A. Velazquez-Campoy. 2024. Extending MeCP2 interactome: canonical nucleosomal histones interact with MeCP2. Nucleic Acids Res. 52:3636–3653.

Richter, K., M. Nessling, and P. Lichter. 2007. Experimental evidence for the influence of molecular crowding on nuclear architecture. J Cell Sci. 120:1673–1680.

Strom, A.R., R.J. Biggs, E.J. Banigan, X. Wang, K. Chiu, C. Herman, J. Collado, F. Yue, J.C. Ritland Politz, L.J. Tait, D. Scalzo, A. Telling, M. Groudine, C.P. Brangwynne, J.F. Marko, and A.D. Stephens. 2021. HP1α is a chromatin crosslinker that controls nuclear and mitotic chromosome mechanics. Elife. 10.

Subramanian, A., P. Tamayo, V.K. Mootha, S. Mukherjee, B.L. Ebert, M.A. Gillette, A. Paulovich, S.L. Pomeroy, T.R. Golub, E.S. Lander, and J.P. Mesirov. 2005. Gene set enrichment analysis: a knowledge-based approach for interpreting genome-wide expression profiles. Proc Natl Acad Sci U S A. 102:15545–15550.

Tanaka, T.U., and A. Desai. 2008. Kinetochore-microtubule interactions: the means to the end. Curr Opin Cell Biol. 20:53–63.

Thiemicke, A., and G. Neuert. 2021. Kinetics of osmotic stress regulate a cell fate switch of cell survival. Sci Adv. 7.

Watanabe, K., K. Morishita, X. Zhou, S. Shiizaki, Y. Uchiyama, M. Koike, I. Naguro, and H. Ichijo. 2021. Cells recognize osmotic stress through liquid-liquid phase separation lubricated with poly(ADP-ribose). Nat Commun. 12:1353.

Xu, P., J. Mahamid, M. Dombrowski, W. Baumeister, A.L. Olins, and D.E. Olins. 2021. Interphase epichromatin: last refuge for the 30-nm chromatin fiber? Chromosoma. 130:91–102.

Zhou, B.R., K.N.S. Yadav, M. Borgnia, J. Hong, B. Cao, A.L. Olins, D.E. Olins, Y. Bai, and P. Zhang. 2019. Atomic resolution cryo-EM structure of a native-like CENP-A nucleosome aided by an antibody fragment. Nat Commun. 10:2301.

